# Chemically induced senescence prompts functional changes in human microglia-like cells

**DOI:** 10.1101/2024.02.27.582168

**Authors:** S. Armanville, C. Tocco, Z. Haj Mohamad, D. Clarke, R. Robitaille, J. Drouin-Ouellet

## Abstract

In response to various stressors, cells can enter a state called cellular senescence which is characterized by irreversible cell cycle arrest and a senescence-associated secretory phenotype (SASP). The progressive accumulation of senescent glial cells in the central nervous system (CNS) with aging suggests a potential role for senescence as driver of aging and inflammation in the brain. As the main immune cell population residing in the CNS, microglia are thought to play a pivotal role in the progression of age-associated neuroinflammation. Furthermore, due to their slow turnover, microglia are highly susceptible to undergoing cellular senescence. However, current understanding of age-related changes in microglia and their impact on brain aging is limited. Due to the challenge in accessing human primary microglia and the lack of models to adequately recapitulate aging, this knowledge is predominantly limited to rodent studies. Here, we chemically induced senescence in a human immortalized microglia cell line with a cocktail of senescence-inducing molecules. We demonstrate that chemically induced senescent microglia adopt a pro-inflammatory phenotype, have reduced phagocytic activity and impaired calcium activity. Our results show that chemically induced senescence can mimic features of cellular aging and can provide insight on the impact of aging and cellular senescence on human microglia.

## Introduction

A growing number of studies suggests that cellular senescence, primarily characterized by permanent cell cycle arrest, is involved in cellular aging. In healthy individuals, senescent cells play a crucial role in wound healing, tissue repair and tumor suppression (1). However, progressive accumulation of senescent cells also correlates with the advancement of age (2), suggesting that this biological process could be a driver of aging. This hypothesis has prompted the development of genetic and pharmacological strategies to eliminate senescent cells in an effort to revert signs of aging in the surrounding environment (3). For instance, the depletion of cells expressing the senescence marker p16^INK4a^ in the brain of older mice delays loss of cognitive function (4). Unlike apoptotic cells, senescent cells maintain an active metabolic state, known as senescence-associated secretory phenotype (SASP). Cells in this state are characterized by gene expression modifications and secretion of pro-inflammatory cytokines that in turn activate the immune system and allow the clearance of senescent cells (5).

In the brain, accumulation of senescent cells during aging suggests their potential implication in neuroinflammation and the onset and progression of age-related neurodegenerative disorders. In the healthy brain, microglia are the resident immune cells and are essential for proper brain functioning and homeostasis (6). However, microglia display a dystrophic morphology in postmortem brain samples from elderly donors (7–9). Additionally, in vivo and in vitro studies show that mouse microglia have defective functionality, including reduced surveillance and phagocytic activity, increased production of reactive oxygen species (ROS) (10, 11), reduced response to insults (12), slower migration to sites of infection and processes shrinkage (13). Further, microglia in old mouse brains show more sustained inflammatory response, which correlates with the highly proinflammatory microenvironment characteristic of the aging brain (11). The release of neurotoxic factors, as well as reduced ability to phagocytize cell debris and toxic protein aggregates, create a potentially harmful environment to neurons (10, 14). However, the causes of age-related microglial dysfunction are poorly understood. Recent studies on the aging mouse brain have reported the existence of p16^high^ microglia, suggesting senescence could play a role in age-related microglial dysfunction (15–17).

In humans, dystrophic microglia have been reported in postmortem brain tissues of patients affected by various neurodegenerative diseases. They have been detected near accumulation sites of the tau protein and beta-amyloid plaques in Alzheimer’s disease (AD), near Lewy bodies in cases of dementia with Lewy bodies (18–20), and a state of intense microglial activation has been observed in mouse models of Parkinson’s disease (PD) and human postmortem samples from PD patients (21, 22). Thus, even though microglial activation is primarily a neuroprotective defense mechanism, its aberrantly sustained activation is suspected to contribute to neuroinflammation in age-related neurodegenerative disorders (23).

To date, only a handful of studies has been conducted on aged human microglia (9, 13, 24, 25). This is likely due to the limited accessibility of these cells, and the lack of non-invasive imaging tools that would allow the study of living aged microglia at the molecular level (26). Therefore, our current understanding of age-related phenotypic changes in microglia stems mainly from studies carried out in rodents, as well as in human postmortem brains which do not allow mechanistic studies. However, inter-species differences in lifespans, phenotypes and age-related changes in microglial function may limit the translatability of rodent findings to human microglia and their contribution to brain aging and neurodegenerative diseases (13, 27, 28).

To dissect the impact of aging and cellular senescence on human microglia, we report a model in which cellular senescence can be induced in microglia-like cells via the exposure to three chemical compounds targeting different homeostatic pathways. SBI-0206965, a known inhibitor of the autophagy initiator kinase ULK1, leads to dysfunctional autophagosomes and affects clearing of damaged mitochondria (29). Additionally, it could lead to aggregation of oxidized proteins, carbonyls proteins and products of lipidic peroxidation and protein glycation found in lipofuscin granules, which are also observed in the cytoplasm of senescent cells (30, 31). Next, the molecule O-151 is an inhibitor of the 8-oxoguanine DNA-glycosylase, a key component of the major pathway implicated in the repair of oxidative DNA damage (32). Previous evidence demonstrates that efficiency of DNA repair systems declines with age which could explain the accumulation of age-associated DNA damage (33–35). Finally, Lopinavir, an HIV protease 1 inhibitor, could impact cellular senescence through its role in disrupting Lamin A processing and leading to progeria-like features in treated cells (36).

Although several strategies to induce senescence in human cells are available, including replication exhaustion, the use of gamma radiation, treatment with genotoxic drugs, induction of oxidative stress and epigenetics modifying agents (37), the use of chemical compounds presents some benefits. For instance, it is more time-efficient compared to other methods such as replicative stress-induced senescence (28), a crucial factor considering the slow turnover of microglia. Furthermore, the use of a combination of small molecules allows targeting several pathways relevant to the ageing of microglial cells at once, including autophagy and DNA repair impairments as well as nuclear lamina alterations, whereas non-chemical methods such as replicative stress and irradiation tend be restricted to targeting one aging process.

In this paper, we show that microglia exposed to the SLO cocktail exhibit classical markers of cellular senescence, adopt a pro-inflammatory phenotype, have reduced phagocytic activity and impaired calcium activity, compared to non-treated cells.

### Experimental procedures

#### Cell line and culture

Immortalized human microglial HMC3 cells were cultured in Eagle’s Minimum Essential Medium (EMEM) supplemented with 10% fetal bovine serum (FBS) and 100U/mL of penicillin/streptomycin solution (P/S) at 37°C and in 5% CO. When cells reached 80-90% confluence, they were trypsinized by incubation with Trypsin-EDTA for 5 min, then collected in fresh complete medium. After centrifuging the cell suspension at 400G for 5 min at room temperature (RT), the supernatant was discarded, and the remaining pellet resuspended in culture medium. Viable cell number was assessed using a Hausser Scientific™ Bright Line™ Counting Chamber and cells were seeded on plastic multi-well culture plates coated with Gelatin 0.1% for 1h at 37°C. For all material and reagent catalog numbers, refer to Suppl. Table 1.

#### Chemical induction of senescence

The chemical compounds SBI-0206965, O-151 and Lopinavir (38) were prepared in dimethyl sulfoxide (DMSO) to obtain a stock concentration of 10 mM, 1 mM and 1 mM, respectively and stored at −80° C. HMC3 cells were seeded at a density of 2,630 cells/cm^2^ on 6, 12, or 24-well plates. After 24h from seeding, cells were treated for 6 consecutive days with a cocktail of SBI-0206965 10 μM, Lopinavir 1 μM, O-151 1 μM or with an equivalent concentration of DMSO in standard HMC3 medium for the full dose treatment, and SBI-0206965 5 μM, Lopinavir 0,5 μM and O-151 0,5 μM for the half dose. On day 6, cells were detached by using trypsin (5 minutes at 37°C) and seeded at a density of 14,285 cells/cm^2^ in 12-well plates to assess phagocytic activity, 8,500 cells/cm^2^ in 8-well, glass-bottomed microchambers for calcium imaging, 45,450 cells/cm^2^ in 48-well plates for inflammatory stimulation or 2,725 cells/cm^2^ for immunostaining analyses.

#### RT-qPCR and bulk RNA-seq

HMC3 cells were collected and centrifuged at 400G for 5 min at RT. The supernatant was discarded, and samples were kept at −80°C until further processing. RNA extraction was carried out following the RNeasy MicroKit protocol and RNA concentration measured using a Nanodrop spectrophotometer. For RT-qPCR, cDNA was synthesized from the extracted RNA using the SuperScriptTM VILO^TM^ Mastermix. ACTB and HPRT1 genes were used as housekeeping genes. The relative expression of genes of interest was determined by quantitative polymerase chain reaction (qPCR) using PowerUp^TM^ SYBR^TM^ Green Master Mix. Data were quantified using the ΔΔCT method. For a list of primers used in this study, refer to Suppl. Table 3. RNA-seq data from HMC3 cells treated with vehicle control (VEH), SLO half dose (SLO 1/2) and SLO full dose (SLO 1) have been deposited at GEO and are publicly available (GSE275256). The dataset was processed and aligned by the IRIC genomic platform at the University of Montreal. Sequences were trimmed for sequencing adapters and low quality 3’ bases using Trimmomatic version 0.38 (39) and aligned to the reference human genome version GRCh38 using STAR version 2.7.1a (40). The quality of the samples was assessed with MultiQC, a report is available as Supplementary Material (Suppl. File 1). Gene expressions were obtained as readcount directly from STAR to obtain normalized gene and transcript level expression. Differential analysis was performed using DESeq2 R package. Gene Set Enrichment Analysis (GSEA) was performed using g:Profiler (41). Further supplementary information is available as supplementary files, including raw reads and mapping statistics (Suppl. File 2), list of differentially expressed genes (Suppl. File 3) and GSEA enriched terms (Suppl. File 4).

#### β-Galactosidase activity

After fixation in 4% paraformaldehyde (PFA) for 15 min, β-Galactosidase (β-Gal) activity was assessed using the CellEvent^TM^ Senescence Green Detection Kit, according to manufacturer’s instructions. Briefly, HMC3 cells were incubated in a solution containing the β-Gal probe diluted 1:1000 in the provided buffer, for 3h in the dark at 37°C. Fluorescent images of 1665px by 1665px were acquired with a Nikon AX/AX R with NSPARC confocal microscope equipped with a 20x lens (PLAN APO 20x DIC M/N2). Positive cells were detected using NIS-Elements analysis software based on β-Gal fluorescence intensity. The percentage of β-Gal positive cells was then calculated by dividing the number of β-Gal positive cells over the total number of cells, identified using 4’,6-diamidino-2-phenylindole (DAPI) nuclear labeling.

#### TUNEL ASSAY

After fixing cells as previously mentioned, apoptosis was assessed using the Click-IT TUNEL Alexa Fluor^TM^ Imaging Assay, according to the manufacturer’s instructions. Briefly, cells were incubated with an enzyme that incorporates modified deoxyuridine triphosphate to 3’-OH ends of fragmented DNA and the fluorescent detection is based on a click reaction that allows to identify fragmented DNA, a hallmark of apoptosis. DNase I, provided in the kit, was used to induce strand breaks in the DNA and was used as a positive control for the TUNEL reaction.

#### Immunocytochemistry

After fixation as previously mentioned, cells were permeabilized and blocked in a solution containing 0.2% Triton-X-100 and 3% bovine serum albumin (BSA) in 0.1M PBS for 1h. Primary antibodies (Suppl. Table 2) were diluted in permeabilization/blocking solution and applied overnight at 4°C. After PBS 1X rinsing, samples were incubated with Fluorophore-conjugated secondary antibodies and DAPI (1:1000), previously diluted in permeabilization/blocking solution, for 2h, at RT and protected from light. Then, samples were rinsed 3 times with PBS 1X prior to imaging. Fluorescent images of 1665px by 1665px and 356px by 356px were acquired with a Nikon AX/AX R with NSPARC confocal microscope equipped with a 20x lens (PLAN APO 20x DIC M/N2). The nuclear area, defined as the DAPI-stained region, was determined using the NIS-Elements analysis software (Nikon) and represented in px^2^. Number of γH2AX foci per cell, was measured as number of γH2AX-positive spots detected within each DAPI-stained region. Number of cytoplasmic chromatin fragments (CCFs), determined by γH2AX and H3K9me3 double positive foci per cell, was automatically counted using NIS-Elements analysis software (Nikon). Briefly, the total number of cells was estimated as number of DAPI-stained nuclei; then, intensity thresholds were set for every staining for object segmentation and to discriminate between positive and negative objects. Finally, the number of CCFs was measured as number of γH2AX/H3K9me3 double-positive spots detected within each phalloidin-stained region and excluded from the DAPI-stained region. For the quantification of the number of β-Gal^+^, pRPS6^+^, pHH3^-^ cells, the total number of cells was estimated as number of DAPI-stained nuclei; then, intensity thresholds were set for every staining for object segmentation and to discriminate between positive and negative objects. The number of single, double and triple positive cells was calculated with the NIS in-built General Analysis tool.

#### Pro-inflammatory stimulation and cytokine secretion analysis

24h after plating, cells were stimulated with a solution of lipopolysaccharide (LPS; 100 ng/mL) and interferon-γ (IFNγ; 10 ng/mL) in standard HMC3 medium for 6h (42). Then, culture medium was collected for ELISA, whereas adherent cells were either processed for RNA extraction and further analyzed by qPCR or fixed for ICC analysis. Post stimulation media were further supplemented with protease inhibitors, centrifuged at 300G for 5 min at 4°C to remove cellular debris and processed as per manufacturer’s instructions to assess secretion of cytokines interleukin 6 (IL-6) and 8 (IL-8), as well as tumor necrosis factor α (TNFα) and interleukin IL-1β. Absorbance was read at 450 nm using a SYNERGY H1 microplate reader, then corrected using the average absorbance values read at 540 and 570 nm.

#### Assessment of phagocytotic capacity by flow cytometry

24h after reseeding, HMC3 cells were incubated with 0.75μm fluorescent carboxylate microspheres for 4h in standard HMC3 medium. Phagocytic capacity was then determined using the CytoFLEX Flow Cytometer instrument and measured as percentage of YFP positive cells. Cells not exposed to beads served as a control to determine the target population based on size. Analysis was performed with the CytExpert software.

#### Calcium imaging

24h after reseeding, HMC3 cells were loaded for 1h with the membrane-permeant Ca^2+^ probe Fluo-4 AM by replacing standard HMC3 medium with loading medium (1 μM Fluo-4 AM, 0.02% Pluronic acid and 1% DMSO, dissolved in standard HMC3 medium). Then, cells were rinsed twice in PBS 1X to remove any Fluo-4 AM residual and left to rest in standard medium condition for 20 min prior to recording. Recording was then carried out in PBS 1X with Ca^2+^ and Mg^2+^, without phenol red. Fluorescent images of 2048px by 2048px (1746 µm by 1746 µm) fields of view were acquired with a Nikon AX/AX R with NSPARC confocal microscope equipped with a 10x lens (PLAN APO λD 10x OFN25 DIC N1). 360 frames were recorded at 1Hz. Specifically, the initial 120 frames before ATP administration were used to define baseline activity, whereas the following 240 frames recorded calcium dynamics elicited by 100nM ATP administration. NIS-Elements software was used to segment single cells and extract signal intensity traces for each individual region of interest (ROI). Each ROI trace was normalized by subtracting background signal intensity values and by dividing by the relative average baseline signal intensity. Among all recorded cells (N= 433 and N= 83 for control VEH and SLO treated conditions, respectively), only cells showing a fold-change at maximum peak of at least 2 from baseline (N = 156; 36% and N= 58; 70% for control VEH and SLO treated conditions, respectively) were included in the analyses. Extrapolation of 90%/10% decay time, peak time, peak amplitude, and area under the curve (AUC) measurements was performed with ClampFit 10.7 open-source software. For time-lapse of microglial calcium imaging, refer to Suppl. videos S1 and S2.

#### Statistical analyses

GraphPad Prism 10 software was used for statistical analysis. Shapiro-Wilk normality test was used to assess the normality of the distribution. When normality could not be assumed, a nonparametric test was performed. Groups were compared using either Bonferroni’s one-way multiple comparison ANOVA, Kruskal-Wallis test with Dunn’s multiple comparisons tests or Bonferroni’s two-way multiple comparison ANOVA. Unpaired Student’s t-test was used in the case of a two-group comparison. An alpha level of 0.05 was defined for significance.

## Results

### Exposure to SLO induces dose-dependent transcriptomic changes in human microglial-like cells

With the aim of generating a cellular model to study neurodegenerative disorders, a recent study identified a combination of three small molecules capable of inducing senescence in neurons differentiated from iPSCs (38). These molecules include an autophagy inhibitor (SBI-020695), a DNA glycosylase inhibitor (O-151) and a Lamin A biosynthesis blocker (Lopinavir). In the study, SLO-treated neurons presented age-related transcriptional alterations comparable to aged human brain tissues (38). Here, we generated a model of aging in human microglia-like cells via the administration of SLO molecules to HMC3 cells, an immortalized human microglia cell line. To determine the minimum effective dose, cells where exposed for 6 consecutive days to either the full dose, as reported by Fathi and colleagues (SLO), or a half dose (SLO ½). Control cells were treated with equivalent concentrations of DMSO (vehicle; VEH). First, to broadly characterize the effects of the SLO treatment, we performed RNA-seq to examine differentially expressed genes between VEH control and the two SLO regimens. The Principal Component Analysis (PCA) showed good segregation between both treatments and controls, as well as between SLO and SLO ½ (Fig. 1a). This result was confirmed by the hierarchical clustering of all replicates (Fig. 1b). After the administration of SLO ½ and SLO, 2961 and 4219 genes were upregulated compared to VEH, and 3375 and 4280 were significantly downregulated, suggesting that both regimens were effective in inducing transcriptional changes (Fig. 1c). Interestingly, the vast majority of SLO ½ regulated genes was also differentially regulated upon administration of the full dose, with the latter affecting substantially more genes, implying that a higher dosage is necessary to influence the expression of certain genes. A volcano plot illustrates the distribution of differentially expressed genes between SLO and VEH (Fig. 1d), whereas a heatmap of all differentially expressed genes with Adjusted P-value under 0.001 and a Log_2_ Fold Change above 2 shows a clear distinction between SLO and VEH gene expression (Fig.1e). The analysis of SLO ½ produced similar results (Suppl. Fig 1a,b).

**Figure 1.**
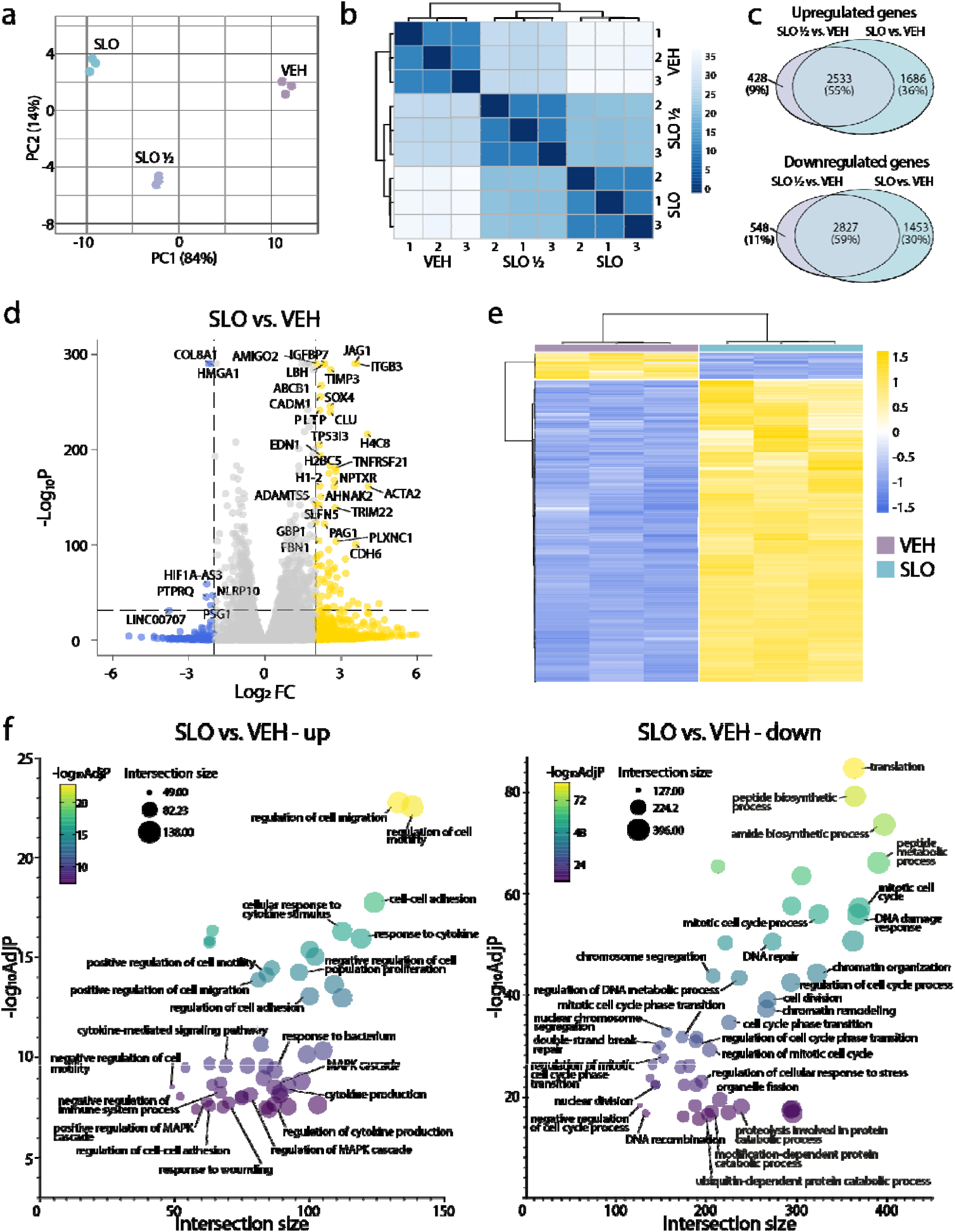
Bulk RNA-seq analysis of HMC3 response to the SLO treatment. (a) PCA plot of vehicle control (VEH), SLO half dose (SLO ½) and SLO full dose (SLO) shows good separation by their top two principal components. (b) Hierarchical clustering of VEH, SLO ½ and SLO representing Euclidean distances among samples. (c) Venn diagrams of upregulated (top) and downregulated (bottom) in SLO ½ vs VEH and SLO vs. VEH comparisons. The intersections represent common differentially regulated genes among conditions. (d) Volcano plot illustrating differentially regulated genes in SLO vs VEH (Log_2_FC > 2 and -Log_10_P > 32). (e) Heatmap of differentially expressed genes in SLO vs VEH (Log_2_FC > 2 and adjusted P-value < 0.001). (f) Bubble plot showing the top 50 enriched Gene Ontology (GO) terms for upregulated (left) or downregulated (right) genes in SLO vs VEH. Further information can be found in Suppl. Files 1-4.

Moreover, a Gene Set Enrichment Analysis (GSEA) of SLO compared to VEH shows that many key biological processes involved in aging were affected by the administration of the SLO treatment. This is illustrated by bubble plots showing the top 50 most enriched terms of upregulated (left) and downregulated (right) genes when comparing SLO full dose to VEH controls (Fig. 1f). Gene ontology (GO) terms either enriched or downregulated in SLO compared to VEH confirmed that SLO treatment in microglia-like cells is effective at blocking the three pathways of interest, serving as confirmation of the treatment efficacy. More specifically, downregulation of autophagy related pathways such as “peptide biosynthetic process”, “proteolysis involved in protein catabolic process”, “ubiquitin-dependent protein catabolic process”, and “modification-dependent macromolecule catabolic process” could be attributed to the ULK1 inhibitor, SBI-020695. Lopinavir, the blocker of Lamin A biosynthesis and disruptor of nuclear architecture, and O-151, the selective inhibitor of OGG1 and blocker of DNA damage repair could both be responsible for defective “chromosome organization”, “DNA damage response”, “protein-DNA complex organization”, “chromatin organization”, “chromosome segregation”. Additionally, among the most relevant upregulated pathways, we found “response to cytokine stimulus”, “negative regulation of proliferation”, “regulation of cytokine production”, all of which are processes that are characteristic of senescence. Vice versa, pathways such as “mitotic cell cycle”, “cell cycle phase transition” and “DNA repair” were all downregulated in SLO-treated cells. Similar results were also obtained in SLO ½ treated cells (Suppl. Fig 1c,d). Together, this RNA-seq data shows that SLO treatment in human microglia-like cells induces transcriptomic changes resembling that of an aging profile.

### Exposure to SLO leads to cell cycle arrest and a senescent-like phenotype in microglia

Cell cycle arrest is one of the main characteristics of cellular senescence (43). Consistently, our RNA-Seq data shows that several genes involved in “positive regulation of cell cycle” are downregulated in SLO ½ and SLO compared to VEH (Fig. 2a, top), and several genes involved in “negative regulation of cell population proliferation” are upregulated (Fig. 2a, bottom). In both instances, DEGs were more affected in SLO treated cells compared to SLO ½, implying a dose-dependent regulation of cell cycle. To test our hypothesis, we measured cell proliferation by quantifying the percentage of cells positive for the mitosis marker phosphorylated histone 3 (pHH3). Our data show a significant reduction of proliferation in the SLO full dose, but not SLO ½, compared to the VEH condition (Fig. 2b,c). Another key senescence marker is the increase in β-Gal activity. However, in recent studies its activity was also observed in conditions of persistent stress, unrelated to senescence (44). Additionally, inactivation of its enzymatic activity does not reverse cellular senescence, suggesting that its specificity as a marker of cellular senescence is limited (45). To overcome this problem, we determined senescent cells as cells positive for β-Gal and phosphorylated RPS6 (pRPS6), an indicator of active protein synthesis, but negative for pHH3 (Fig. 2b), following a published algorithm that allows to discriminate among cycling, stressed, quiescent and senescent cells (44). In our system, only microglia treated with the full dose shows an increase in senescent cells percentage (Fig. 2c,d). In a similar fashion, the percentage of p21+ cells, a cell cycle arrest marker, were significantly increased in SLO ½ and SLO treated cells compared to vehicle controls (Fig. 2e,f). This was also the case at the transcript levels when measuring the expression of the CDKN1A gene (Fig. 2g). Interestingly, a clear signature of programmed cell death did not emerge from the RNA-sequencing data. To confirm that the exposure to the small molecules was not causing cell death, we measured cell death-associated DNA fragmentation via a TUNEL assay (Suppl.Fig. 2a), and Draq7 uptake by flow cytometry (Suppl.Fig. 2b). Both assays confirmed that apoptosis was not induced by SLO ½ or SLO, compared to VEH control.

**Figure 2.**
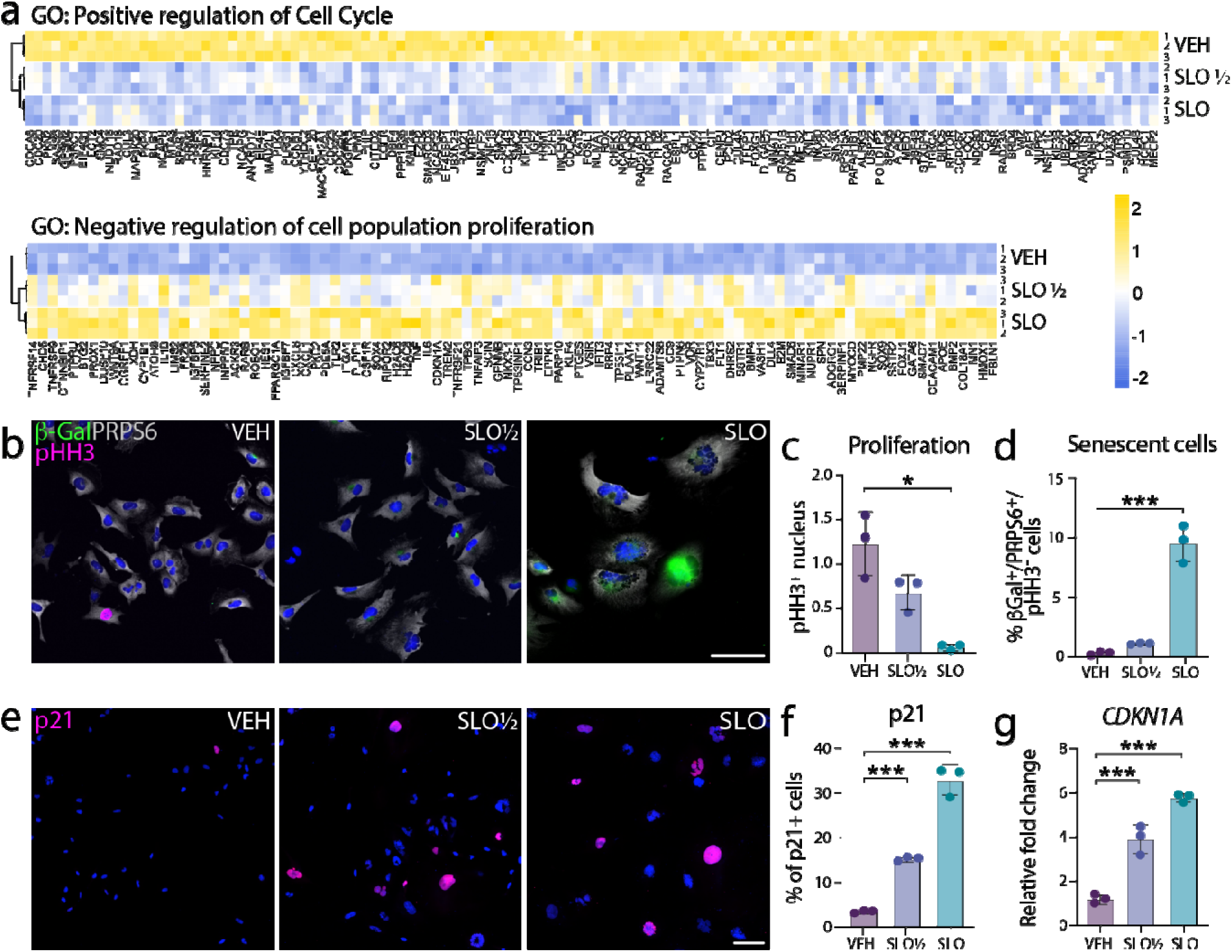
SLO full dose but not half dose promotes cell cycle arrest and features of cellular senescence in HMC3 cells. (a) Expression profile of differentially regulated genes belonging to the Positive regulation of cell cycle (top) and Negative regulation of cell population proliferation (bottom) gene ontology terms. (b) Representative fluorescence images showing microglia treated with VEH, SLO ½ and SLO co-stained with β-Gal, pHH3 and PRPS6. DAPI (blue) is used as nuclear counterstaining. (c) Cell proliferation quantified as the number of pHH3 positive nucleus over the total number of cells. (d) Percentage of senescent cells quantified as the percentage double positive β-Gal and PRPS6 cells but negative for pHH3. (e) Representative fluorescence images showing p21 positive cells in microglia treated with VEH, SLO ½ and SLO. DAPI (blue) is used as nuclear counterstaining. (f) Quantification of percentage of p21 positive cells over total number of cells. (g) RT-qPCR quantification of relative gene expression of the p21 coding gene CDKN1A in microglia treated with VEH, SLO ½ or SLO. Data for all graphs are represented as mean ± SD. [*p<0.05; **p<0.01; ***p<0.001 by one-way ANOVA, n=3 independent experiments]. Scale bar = 100 μm (low magnification) and 20 μm (high magnification).

### Exposure to SLO leads to nuclear morphological and architectural changes characteristic of cellular senescence

During cellular senescence, specific changes in nuclear morphology and architecture can occur. Interestingly, loss of Lamin B1, an important structural component of the nuclear lamina implicated in modulation of nuclear functions and whose expression decreases in senescent cells, is observed upon SLO administration (Fig. 3a,b) (46). Senescent-associated morphological changes such as an increase in nucleus area, structural disruption, invaginations and enlargement or fragmentation of the nucleus (47–50) were observed in SLO treated cells (Fig. 3a,c). As a result of nuclear alterations, chromatin fragments can escape the nucleus and accumulate in the cytoplasm as CCFs and constitute another hallmark of cellular senescence. These CCFs are usually enriched in H3K9me3 positive heterochromatin and are positive for the DNA damage marker γH2AX (49). We found that H3K9me3 and γH2AX double-positive fragments were increased in the cytoplasm of SLO treated cells, compared to VEH controls (Fig. 3d,e). Finally, since senescent and aged cells are reported to accumulate double-strand DNA damage (34, 51, 52), we next assessed whether SLO could induce such accumulation in HMC3 cells. For this purpose, we quantified the number of γH2AX foci per nucleus and observed that SLO full dose, but not half dose, significantly induces double-strand DNA damage, compared to vehicle controls (Fig. 3d,f).

**Figure 3.**
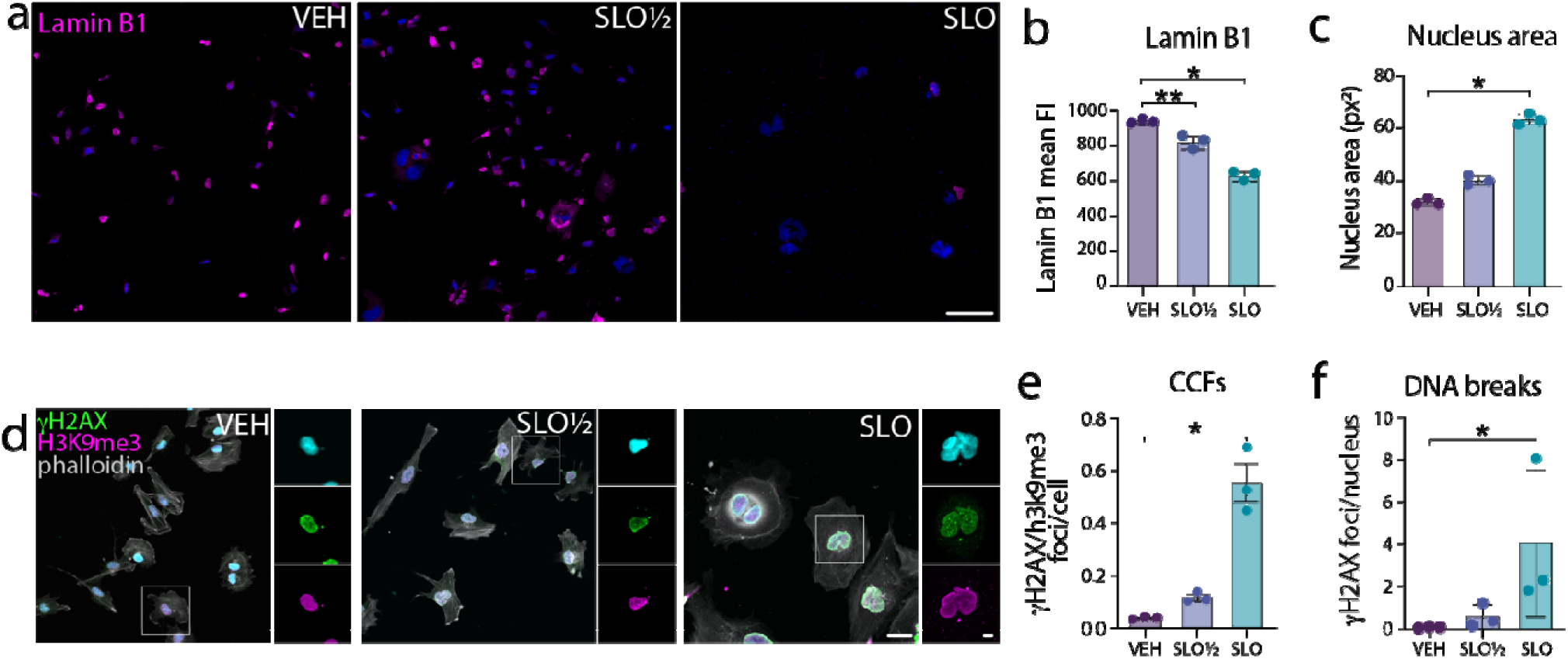
SLO full dose but not half dose induces changes in nuclear morphology and architecture characteristic of cellular senescence. (a) Representative fluorescence images showing microglia treated with VEH, SLO ½ and SLO stained with Lamin B1. DAPI (cyan) is used as nuclear counterstaining. (b) Quantification of Lamin B1 fluorescence intensity in the nucleus. (c) Nucleus area determined by core surface area (px). Each point corresponds to the mean ± standard deviation of the core area of 15 microglia. (d) Representative fluorescence images showing microglia treated with VEH, SLO ½ and SLO co-stained with γH2AX, H3K9me3 and phalloidin. DAPI (blue) is used as nuclear counterstaining. (e) Number of cellular cytoplasmic chromatin fragments determined as the number of γH2AX/H3K9me3-positive foci per cell in the cytoplasm only. (f) Number of DNA double-strand breaks measured as number of γH2AX-positive foci per nucleus delineated by DAPI. Each point corresponds to the mean ± the standard deviation of the number of γH2AX foci per nucleus of 15 microglia. Data for all graphs are represented as mean ± SD. [*p<0.05; **p<0.01; ***p<0.001 by one-way ANOVA, n=3 independent experiments]. Scale bar = 100 μm (low magnification) and 20 μm (magnification).

### Exposure to SLO exacerbates the pro-inflammatory response

With age, microglia adopt a “primed” phenotype and homeostatic function is thought to become increasingly disrupted over time, a process that is deemed to be implicated in the progression of neurodegenerative diseases (10, 23, 53). Interestingly, SLO treated microglia also showed an enrichment of genes associated with neurodegenerative disorders (Suppl.Fig 3a). Consistently, our GSEA analysis shows enrichment in multiple signatures of microglia activation, including cytokine secretion and response to cytokines for both SLO ½ and SLO, although expression differences are more pronounced upon full SLO dose (Fig. 4a). Since we could not detect strong molecular and structural changes upon SLO ½ administration, we decided to focus on the comparison between VEH controls and SLO full dose for the functional characterization of treated microglia. Primed microglia adopt an overactive phenotype by secreting an excessive amount of pro-inflammatory cytokines that can be further exacerbated after a pro-inflammatory stimulation, like LPS (54). To confirm that the overexpressed cytokines were effectively secreted upon SLO stimulation, and that this proinflammatory profile could be exacerbated by the administration of LPS and Interferon-γ, we exposed VEH- and SLO-treated HMC3 for 6 hours to either a combination of the proinflammatory stimuli LPS and IFNγ (LPS/IFNγ), or PBS (CTL). After the proinflammatory stimulation, gene expression of common inflammatory cytokines IL-1β, IL-6, IL-8 and TNF was assessed by RT-qPCR and protein secretion in the supernatant by ELISA. In line with our hypothesis, the transcript expression of all cytokines was significantly increased upon pro-inflammatory stimulation in SLO-treated microglia compared to control vehicle. Among them, IL-1β was the only cytokine to show a higher relative expression already at basal levels (Fig. 4b). Additionally, both IL-6 and IL-8 showed increased secretion post-stimulation in SLO treated cells, compared to control (Fig. 4c), but secretion of IL-1β and TNFα was not detected in any of the conditions. According to literature, HMC3 cells are unable to secrete TNFα (55) whereas the expression of IL-1β has been verified at both mRNA and protein level by western blot, but its secretion in the media remains elusive, possibly due to high turnover rate of the mature protein (56, 57). Importantly, secretion of both IL-6 and IL-8 was already higher at baseline in SLO-treated cells compared to controls, in line with the senescent-like profile exhibited by these cells. It has been reported that, in senescent cells, chromatin associated non-histone binding protein HMGB1 (58), which normally localizes in the nucleus, is translocated to the cytosol and then secreted extracellularly (59), where it acts as a damage-associated molecular pattern (DAMP) and induces inflammation to the site of tissue damage (60, 61). In the extracellular space, HMGB1 modulates senescence by binding to the surface of immune cells to initiate the signaling pathway responsible for the expression of pro-inflammatory cytokines (60). We observed a significant decrease in HMGB1 expression in the nucleus, accompanied by an increase in secreted HMGB1 (Fig. 4d-f), suggesting that it was mostly secreted to mediate the SASP. Together, our results show that SLO treated cells acquire pro-inflammatory features, including secretion of key pro-inflammatory cytokines, and that this process could be HMGB1 dependent.

**Figure 4.**
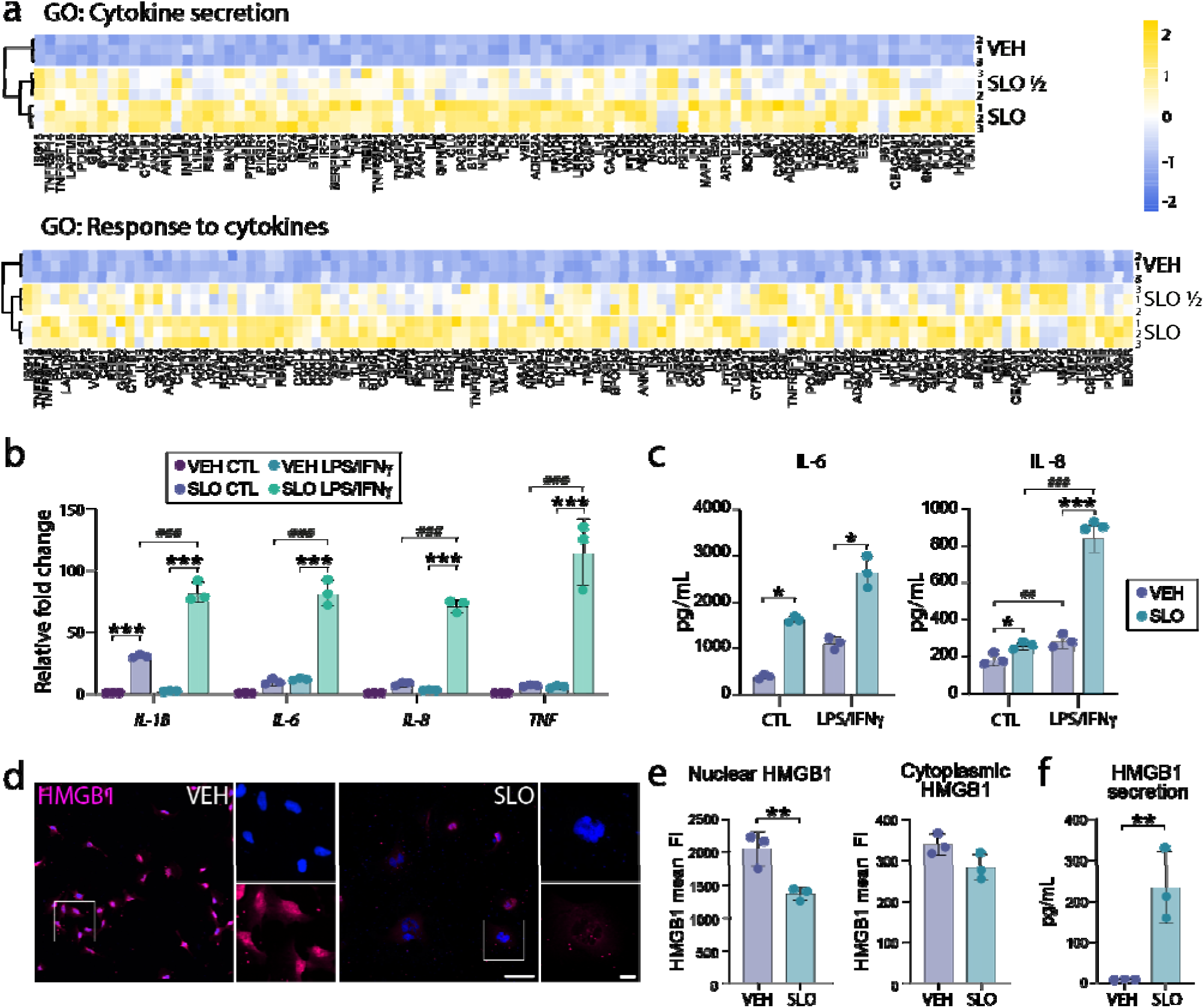
SLO treated HMC3 show an overly reactive, pro-inflammatory profile. (a) Expression profile of differentially regulated genes belonging to the cytokine secretion (top) and Negative regulation of response to cytokines (bottom) gene ontology terms. (b) RT-qPCR quantification of relative gene expression of SASP-associated genes (IL-1B, IL-6, IL-8, TNF) in microglia treated with VEH or SLO either stimulated with pro-inflammatory stimulation (LPS/IFNγ) or untreated (CTL). (c) Protein concentration (pg/mL) of IL-6 and IL-8 in culture medium from microglia treated with vehicle (VEH) or senescence molecules (SLO) either stimulated with pro-inflammatory molecules (LPS/IFNγ) or untreated (CTL), measured by ELISA. (d) Representative fluorescence images showing microglia treated with VEH and SLO stained with HMGB1. DAPI (blue) is used as nuclear counterstaining. (e) Quantification of HMGB1 fluorescence intensity in the nucleus and in the cytoplasm. (f) Protein concentrations (pg/mL) of HMGB1 secreted in culture medium from microglia treated with VEH and SLO, measured by ELISA. Data for all graphs are represented as mean ± SD. [*p<0.05; **p<0.01; ***p<0.001 by one-way ANOVA, n=3 independent experiments]. Scale bar = 100 μm (low magnification) and 20 μm (high magnification).

Another signaling pathway that is emerging as master regulator of SASP secretion in response to DNA fragments leakage in the cytosol, is the Stimulation of interferon genes (STING) pathway. Our GSEA analysis pointed to the activation of the “response to viruses” process, another key function of the cGAS-STING pathway, in SLO treated cells, compared to controls (Suppl.Fig 3b). We further validated this finding by measuring transcriptional levels of several cGAS-STING pathway-related molecules and effectors upon SLO stimulation (Suppl.Fig 3c). Interferon β1 (IFN-β1) and C-X-C motif chemokine ligand 10 (CXCR10) were upregulated by over 1000 folds, although this did not reach significance.

Other interferon-stimulated genes such as interferon-induced transmembrane protein 3 (IFITM3), Oligoadenylate Synthetase (OAS) and Chemokine (C-C motif) ligand 5 (CCL5) showed a significant up-regulation, hinting to a cGAS-STING pathway activation. However, further investigation would be required to elucidate whether the activation of this pathway upon SLO administration is responsible to induce SASP secretion in our model.

### SLO-treated microglia shows impairment in fundamental cellular functions

In addition to the pro-inflammatory secretion profile, aged microglia have been reported to show impairment of phagocytic capacities (62–64). To test if SLO exposure caused similar defects, we evaluated phagocytic activity by measuring intake efficiency of fluorescent beads over a 3h period (Fig 5 a,b). Our results show a significant decrease of bead intake in SLO-treated cells compared to VEH-treated cells. Since both phagocytosis and autophagy rely on similar machineries, we have also tested the ULK1 inhibitor SBI-0206965 individually. A similar decrease was observed in SBI-0206965-treated cells, while the two other molecules tested individually did not impact on phagocytic activity, suggesting that blocking autophagy through ULK1 inhibition is necessary and sufficient to induce phagocytic impairment in SLO treated cells (Supp. Fig. 4a,b).

**Figure 5.**
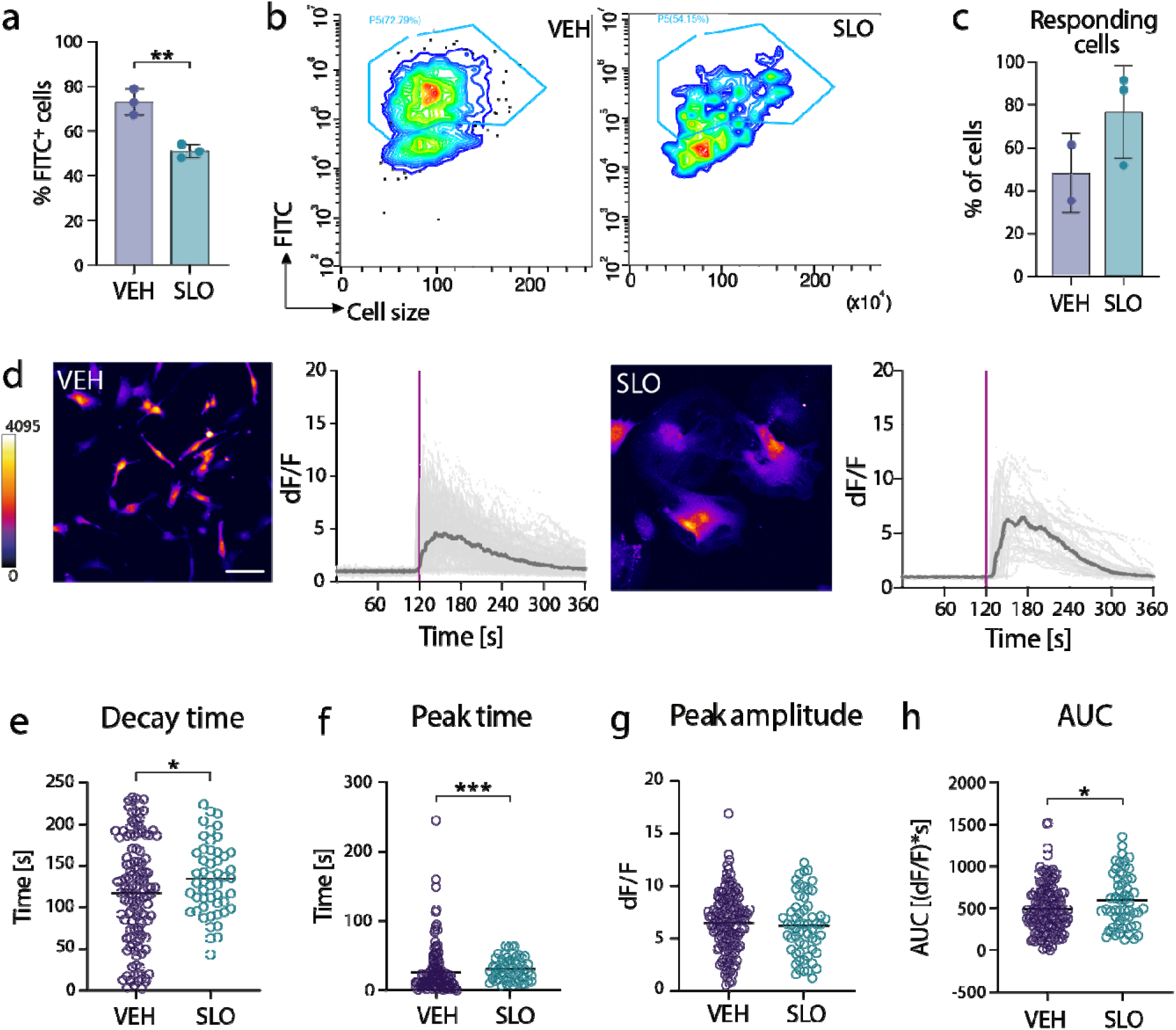
SLO treated microglia show an aberrant functionality. (a) Phagocytosis efficiency determined by the percentage of cells that incorporated fluorescent beads by flow cytometry. (b) Flow cytometry analysis showing the incorporation of fluorescein isothiocyanate-labelled (FITC) beads by microglia after treatment with VEH or SLO. (c) Quantification of the percentage of cells responding to ATP stimulation. Data are presented as mean ± SD. (d) Individual cells (light grey) and average traces (dark grey) Ca responses elicited by administration of 100nM of ATP (purple line) in control vehicle and SLO treated HCM3 cells. Insets show representative MIP confocal images of the Ca level in H cells loaded with 1µM CaC3 indicator Fluo-4 AM. Representative images of the Fluo-4 AM sign l is pseudo colored. (e-h) Scatter plots showing 90%/10% decay time (e), peak time (f), peak amplitude (g), and area under the curve (AUC, h) measurements for n=156 vehicle- and n=58 SLO-treated cells, collected from N=2 (VEH) and N=3 (SLO) technical replicates.. Data for all graphs are represented as mean ± SD. Scale bar = 50μm. [*p<0.05; **p<0.01; ***p<0.001; compared to VEH; #p<0.05; ##p<0.01; ###p<0.001; compared to the CTL group].

Finally, to further characterize the effects of the SLO cocktail on microglial function, we tested whether intracellular calcium (Ca^2+^) dynamics were altered upon SLO administration. In physiological conditions, dynamics of intracellular calcium are known to modulate several properties and functions of microglia, including proliferation and phagocytosis (65). Further, a recent study conducted in the aging mouse brain reported impairment in in vivo Ca^2+^ activity in microglia compared to young and middle-aged brains (66). Upon 30 min incubation with Fluo4-AM fluorescent calcium indicator, cells were recorded over the course of 6 min, including 2 min of baseline activity, and 4 min of activity post ATP stimulation (Fig. 5c). Interestingly, while there was a trend towards a higher percentage of cells responding to ATP stimulation in the SLO condition (Fig. 5d), SLO treated cells depicted slower calcium dynamics, with longer decay time (Fig. 5e), delayed peak time (Fig. 5f) and increased AUC (Fig. 5h) compared to control cells, while peak amplitude remained similar (Fig. 5g). Together, our results show that microglia cells treated with the senescence inducing cocktail SLO acquire features of aging microglia such as increased immunoreactivity, reduced phagocytosis capacities and aberrant calcium dynamics.

## Discussion

The goal of this study was to develop a model that recapitulates well the impact of aging and cellular senescence on human microglial functions. We show that the SLO molecules induce cell cycle arrest in microglia-like cells, a feature that was not assessed in neurons treated with SLO given that neurons are post-mitotic. In our model, an increase in some of the cell cycle arrest markers was also accompanied by senescence-related changes of nuclear morphology and architecture and transcriptional and functional changes, characteristics of aged microglia. Finally, senescent microglia adopted a pro-inflammatory phenotype which was accompanied by aberrant functionality such as reduced phagocytic activity and impaired calcium activity.

In healthy individuals, microglia ensure homeostasis in the central nervous system, by constantly monitoring the presence of endogenous and exogenous harmful substances in their microenvironment. As such, they are in continuous movement to cover long distances and monitor large areas of the brain (67), and exhibit dynamic behavior and ability to quickly switch to an activated state and adopt anti- or pro-inflammatory phenotypes upon detection of danger (67). However, with age, microglial homeostatic functions are gradually disrupted. Possibly due to previous exposure to harmful factors, aging microglia are often reported as being primed or reactive, and can respond in a more robust and sustained manner to an insult (67, 68), while showing reduced phagocytosis activity (69). Similar functional changes are also reported in senescent microglia induced by increased proliferation in a mouse model of AD (70), suggesting cellular senescence could constitute a common source of age-related functional alterations in the context of neurodegenerative diseases. Our transcriptomic data shows that chemically induced senescent microglia are characterized by a signature reminiscent of aging microglia, including differential regulation of many genes related to cell motility, cell cycle arrest, increased production of and response to cytokines. To date, the mechanisms through which senescence affects cellular function remain elusive. It has been proposed that changes in the homeostatic genes expression profile that occurs with age could explain these phenotypical changes (10). Age-related sustained microglial activation could also be explained by the enhancing effect of SASP components, namely IL-6 and IL-8 (71). In this study, both cytokines are released upon SLO treatment. It has been reported that SASP factors can reinforce senescence in both autocrine and paracrine fashion. In this context, the use of co-cultures of microglia and neurons would help to elucidate the role of senescence in brain aging. Some evidence suggests that with age, neurons adopt a senescence-like phenotype and studies conducted in rodents showed that neurons can acquire a SASP-like phenotype and induce premature paracrine senescence in embryonic fibroblasts (72, 73). Co-cultures of neurons and microglia could provide additional insight in this regard.

Analysis of calcium activity in microglia has been used to study their state of activation, as intracellular calcium signaling allows to coordinate and perform microglial executive functions (74). Results from a study conducted in mice revealed age-related changes in intracellular calcium signaling (66). In this paper, the authors identified in middle-aged mice a subpopulation of microglia showing a more reactive phenotype and characterized by increased spontaneous calcium transients. This population of cells was more abundant than in young animals. Interestingly, this phenomenon was further exacerbated in old mice, where another microglia subpopulation characterized by a more senescent or dysfunctional phenotype was reported, with faster migration of microglial processes toward ATP-sources albeit in a more disorganized manner (66). In our work, we show that although SLO-treated cells still display robust calcium activity in response to ATP, they exhibit calcium dynamics distinct from VEH-treated cells. This could have an impact on phagocytosis proficiency, cellular motility, and the metabolic state of SLO-treated microglia. However, a clear link between dysfunctional calcium signaling and microglial functions in our system has yet to be determined.

Nuclear and DNA integrity, as well as autophagy, are essential elements of healthy cells, and their disruption often results in accelerated aging and progression of degenerative diseases. In this paper, we report that perturbation of such mechanisms leads to senescence in human immortalized microglia cells. Furthermore, the three different molecules composing the SLO cocktail not only affect these three key elements at once, but they could also synergize, and cause even deeper disturbances (Fig. 6). For instance, the main function of Lopinavir is to block the production of mature Lamin A via inhibition of the enzyme ZMPSTE24 (36), in turn disrupting the nuclear lamina architecture. However, loss of Lamin A has also been associated to impaired DNA damage repair and increased vulnerability to DNA-damaging agents such as oxidative stress (75). Additionally, O-151 acts as an inhibitor of Oxoguanine glycosylase OGG1, a key component of oxidative DNA damage repair (32). Thus, the two molecules could act in concert to both cause DNA damage and disrupt DNA repair, leading to accumulation of chromatin fragments and their release into the cytoplasm. Furthermore, by blocking autophagy, SBI-020695 leads to accumulation of dysfunctional proteins and organelles (76), including the nucleus, leading to further accumulation of DNA and mitochondria damages, causing an increase in ROS production. While HMGB1 could act as a direct mediator of inflammation in the extracellular space (77), the cGAS-STING signaling pathway could induce transcription of inflammatory cytokines of the SASP via the transcription factors IRF3 and NF-κB, and p21-dependent cell cycle block (78). Further studies are required to dissect the complexity of this network and help clarify the cellular mechanisms triggered by the SLO cocktail.

**Figure 6.**
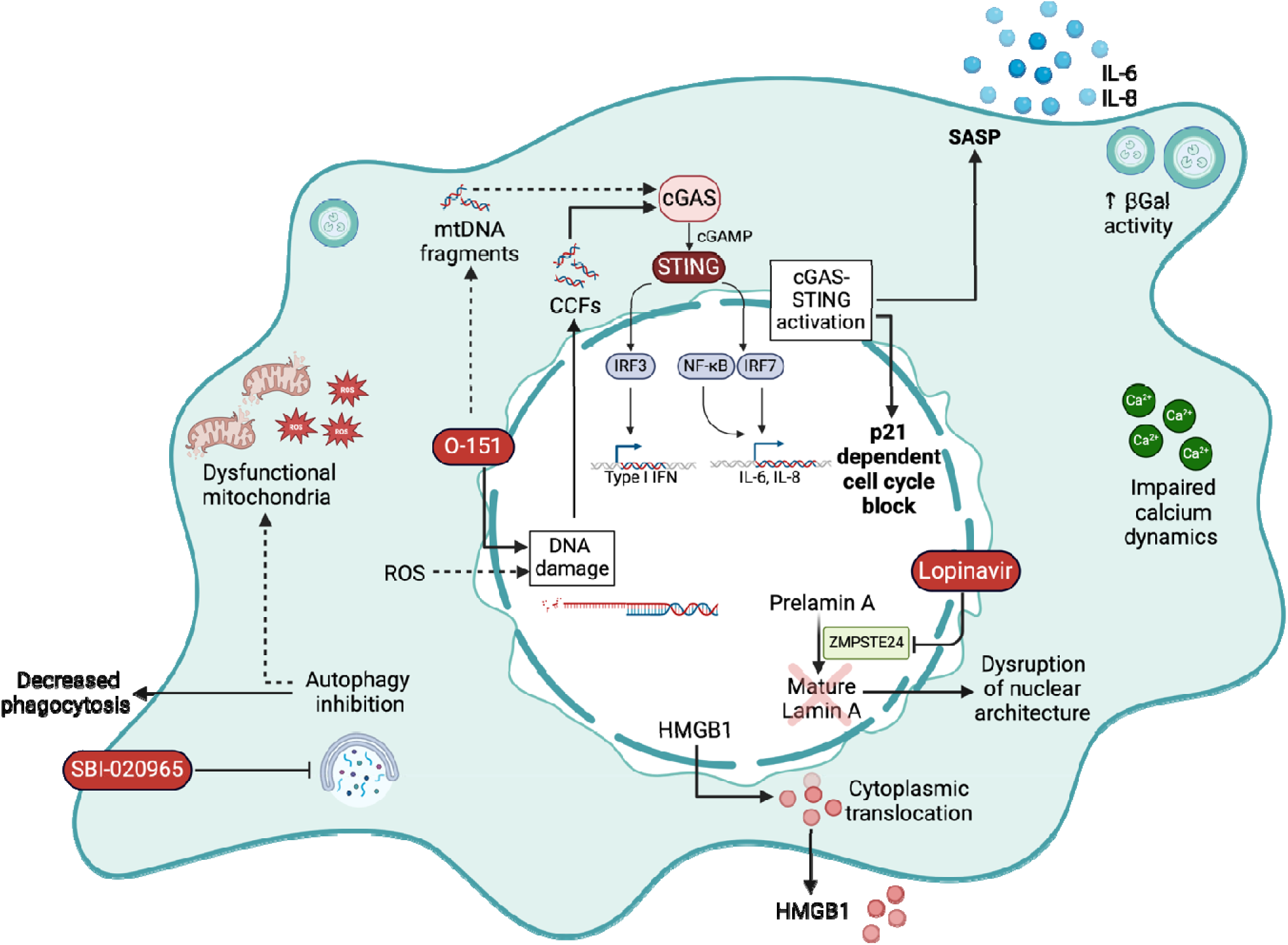
Targeting multiple homeostatic pathways via exposure to SLO mimics the multifactorial aspect of cellular senescence in microglia-like cells. Lopinavir blocks the production of mature Lamin A which disrupts the nuclear lamina architecture and is also associated with impaired DNA damage repair and increased vulnerability to DNA-damaging agents, such as oxidative stress. O-151 blocks oxoguanidine glycosylase OGG1, a key component of oxidative DNA damage repair. The two molecules could synergize, and both disrupt DNA damage repair leading to accumulation of cytoplasmic chromatin fragments (CCFs). As a blocker of autophagy, SBI-020695 leads to accumulation of dysfunctional proteins and organelles, including the nucleus, leading to further accumulation of unrepairable DNA damage and mitochondria, causing an increase in reactive oxygen species (ROS) production. Excessive ROS can lead to further mitochondrial and irreparable nuclear DNA damage. Finally, accumulation of DNA damage can cause a catastrophic chain of events that could culminate in HMGB1 secretion, and/or activation of the cGAS-STING pathway, in response to CCF accumulation and mitochondrial DNA (mtDNA) fragments.

Despite presenting obvious advantages such as easy and reproducible handling and a high rate of proliferation, the use of immortalized cell lines is a limitation to study the effects of aging and senescence on cellular functions. HMC3 cells were generated via immortalization of human microglia by SV40T transfection (79) and as such, we cannot exclude that the method used for immortalizing HMC3 cells, which disrupts the p53 and Rb pathways, interferes with the gene expression of some of the cyclins and other cell-cycle related genes. For instance, the key cell cycle arrest and senescence gene CDKN2A (p16 ^INK4A^) is not upregulated in our system. This suggests that the cells are in a ‘pre-p16^INK4A^ engagement’ phase, as p16^INK4A^ is often induced in the later stages of senescence, whereas p21 is activated much earlier and is sufficient to induce cell cycle arrest. Furthermore, HMC3 cells display lower phagocytic activity than primary microglia (42) and some studies also reported that expression the inflammatory mediators IL-1β, IL-6, TNFα, iNOS, VEGF and TGFβ1 could only be confirmed at the mRNA levels but were not observed at the protein level (42, 80, 81), as we also report to some extent in our work. Finally, the HMC3 immortalization process also altered some original microglial antigenic characteristics, such as the loss of expression at basal level of CD68 and CD11b, two markers of activated microglia (42). Moving forward, the use of iPSC-derived microglia would be a preferred human microglial model, as it more closely resembles primary microglia. However, immortalized cells tend to be easier to culture than iPSC cultures and have been modified to proliferate indefinitely suggesting that SLO concentrations would need to be adjusted to induce cellular senescence while avoiding toxicity and induction of cellular death.

The combination of a renewable source of microglia, with a reliable protocol to induce cellular senescence will help gain insight on the potential role of microglial senescence in the aging brain and the progression of neurodegenerative disorders.

## Acknowledgements

We thank Nicolas Giguère (University of Montreal) for his valuable help with high content imaging, and Gareth Palidwor (University of Ottawa, Bioinformatics Core Facility) for offering his expertise and insights into RNA-seq data analysis. This project has been made possible by the Brain Canada Foundation through the Canada Brain Research Fund with the financial support of Health Canada and the Azrieli Foundation as well as by funding from the Canada Research Chair Program to J.D.O. C.T is receiving a Postdoctoral fellowship from Parkinson Canada. Z.H.M received a Natural Science and Engineering Research Council undergraduate scholarship.

## Author contributions

Sandrine Armanville: conceptualization, methodology, formal analysis, investigation, writing – original draft, review and editing, visualization; Chiara Tocco: conceptualization, methodology, formal analysis, investigation, writing – review and editing, visualization; Zaya Haj Mohamad: investigation, writing – review and editing; Darren Clarke: methodology, writing - review and editing. Richard Robitaille: writing – review and editing; Janelle Drouin-Ouellet: conceptualization, methodology, writing – review and editing, visualization.

## Preprint publication Statement

A preprint of this manuscript has previously been published on BioRxiv (82).

