## Supplementary for "Chemically induced senescence prompts functional changes in human microglia-like cells"

**Supplementary materials**

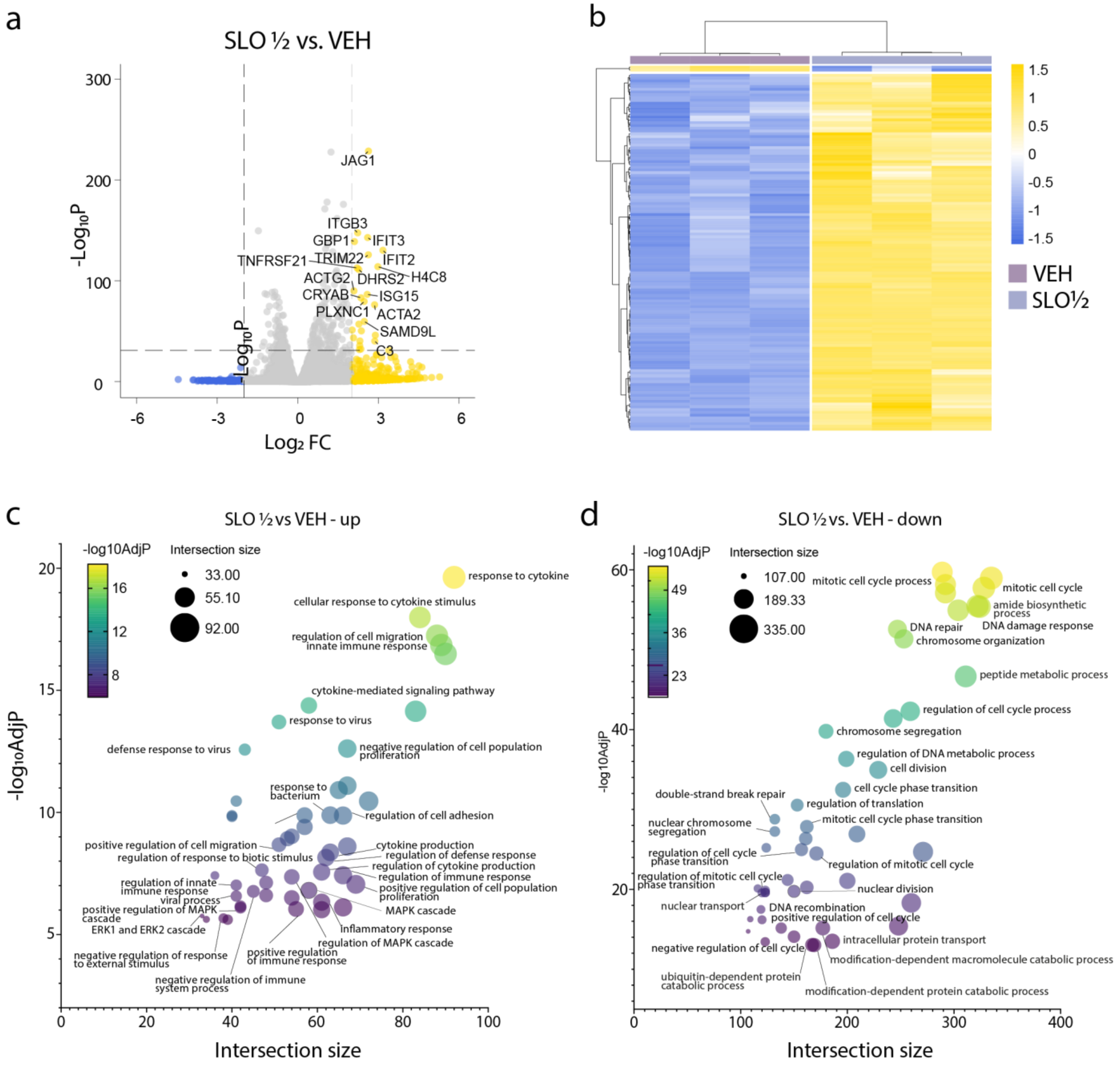


**Supplementary Figure 1. Bulk RNA-Seq transcriptome analysis of HMC3 response to SLO ½ ageing treatment.** (**a**) Volcano plot illustrating differentially regulated genes in SLO ½ vs VEH ($Log_2FC > 2$ and $-Log_{10}P > 32$). (**b**) Heatmap of differentially expressed genes in SLO ½ vs VEH ($Log_2FC > 2$ and adjusted P-value < 0.001). (**c-d**) Bubble plot showing the top 50 enriched Gene Ontology (GO) terms for upregulated (**c**) or downregulated (**d**) genes in SLO ½ vs VEH. (Further information can be found in **Suppl. Files 1-4**).

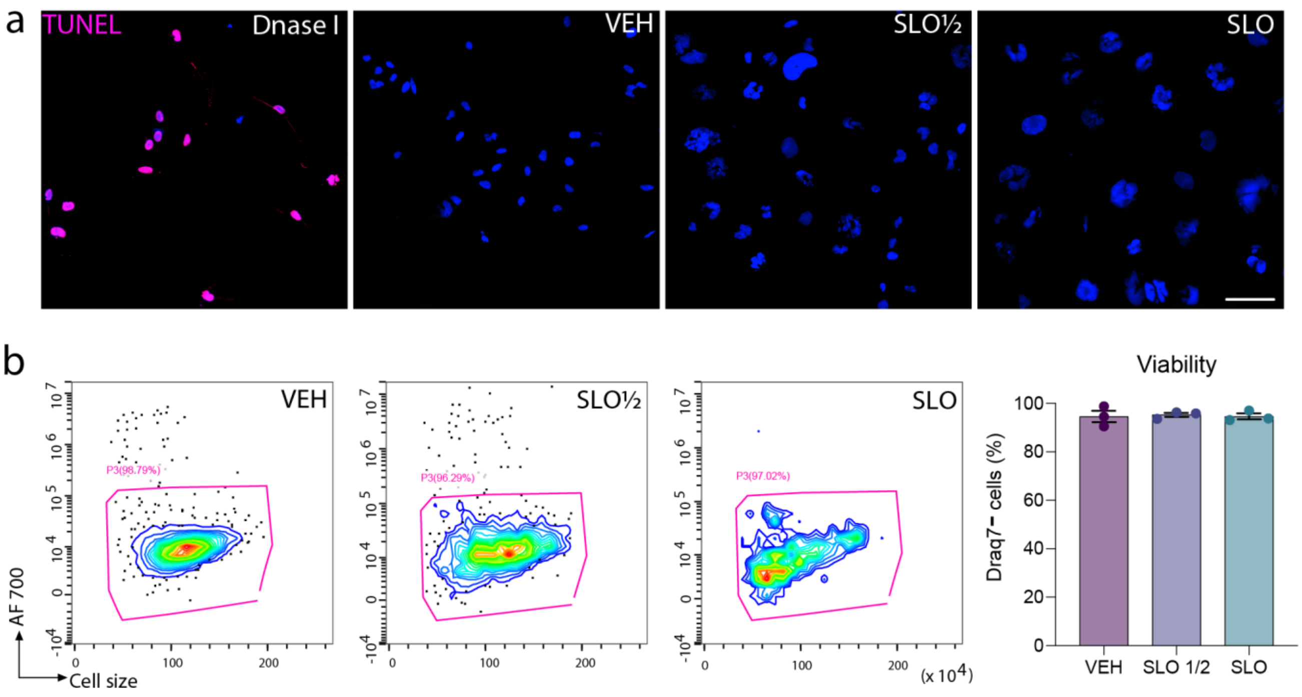


**Supplementary Figure 2. SLO treatment does not trigger cell death in HMC3 cells.** (**a**) Representative fluorescence images showing absence of TUNEL positive cells in VEH, SLO ½ and SLO treated HMC3 cells. HMC3 cells treated with DNase I were used as positive control, as per manufacturer recommendations. (**b**) Flow cytometry analysis of cell viability, shown as incorporation of the Draq7 probe by microglia after treatment with VEH, SLO ½ or SLO. Data are presented as mean ± SEM.

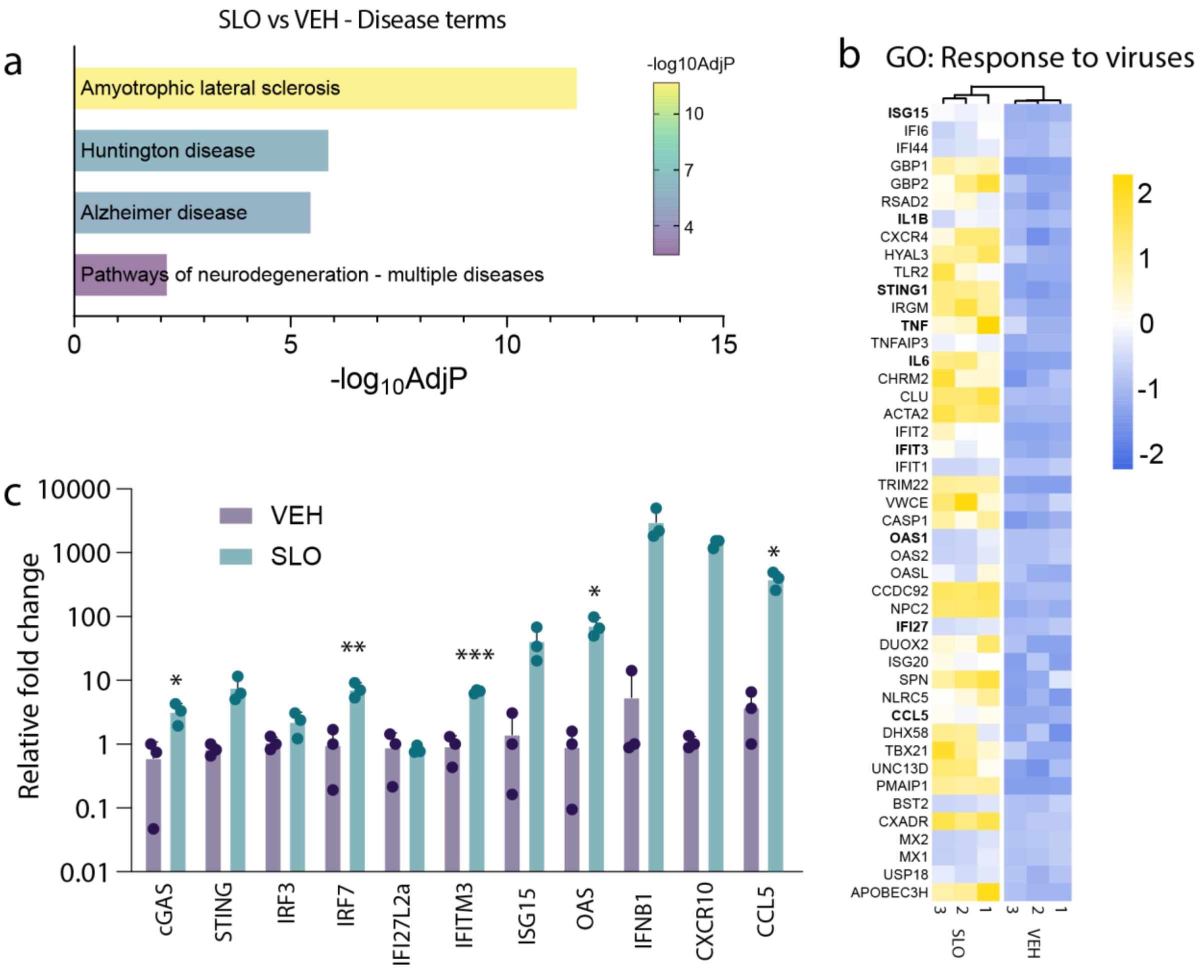


**Supplementary Figure 3. SLO treatment triggers response to viruses and cGAS-STING pathway activation in microglia.** (**a**) Gene Ontology (GO) terms related to Disease for upregulated genes in SLO vs VEH. (**b**) Expression profile of differentially regulated genes belonging to the "Response to viruses" gene ontology term in VEH and SLO treated HMC3. (**c**) RT-qPCR quantification of relative gene expression of cGAS, STING, and several downstream effectors in control VEH and SLO treated microglia.

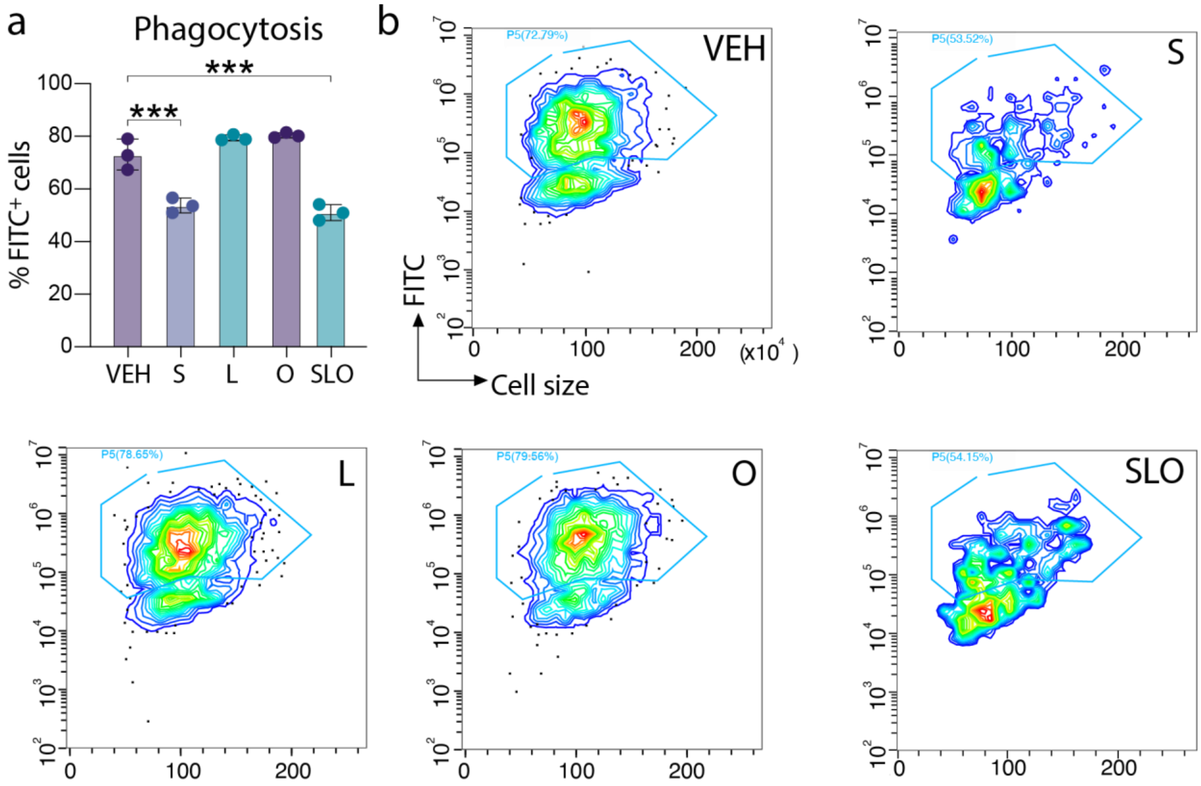


**Supplementary Figure 4. ULK1 inhibitor SBI-0206965 is necessary and sufficient to induce phagocytosis impairment in microglia.** (**a**) Flow cytometry analysis showing the incorporation of fluorescein isothiocyanate-labelled (FITC) beads by microglia after treatment with VEH, SBI-0206965 (S), Lopinavir (L), O-151 (O) or SLO. (**b**) Phagocytosis efficiency is determined by the percentage of cells that incorporated FITC fluorescent beads by flow cytometry.

**Table S1. List of materials and reagents**

| Reagent Name | Company | Catalogue number |
|---|---|---|
| Human embryonic Microglia Clone 3 (HMC3) | ATCC | CRL-3304 |
| Eagle's Minimum Essential Medium (EMEM) | Wisent | 320-026-CL |
| Foetal bovine serum (FBS) | Wisent | 090-150 |
| penicillin/streptomycin solution (P/S) | GIBCO | 15140122 |
| Trypsin-EDTA | GIBCO | 15400054 |
| Hausser Scientific™ Bright Line™ Counting Chamber | ThermoFisher | 02-671-51B |
| Gelatine | Sigma | ES-006-B |
| Dimethyl Sulfoxide (DMSO) | Sigma | D1435 |
| SBI-0206965 | StemCell | 100-0270 |
| O-151 | Carbosynth | BOI168359 |
| Lopinavir | Sigma | SML1222 |
| CellEventTM Senescence Green Detection Kit | ThermoFisher | C10850 |
| 4',6-diamidino-2-phenylindole (DAPI) | Sigma | D9542 |
| PFA | Sigma | P6148 |
| Click-IT TUNEL Alexa Fluor™ Imaging Assay | ThermoFisher | C10247 |
| Triton-X-100 | Sigma | x100 |
| BSA | Sigma | A9418 |
| Donkey serum | Cederlane | 017-000-121 |
| 0.75µm fluorescent carboxylate microspheres | Cederlane | 776610 |
| 8-well, glass-bottomed microchambers | ThermoFisher | 154453 |
| Fluo-4 AM | ThermoFisher | F14202 |
| Pluronic acid | ThermoFisher | P6867 |
| Adenosine 5'-triphosphate disodium salt hydrate (ATP) | Sigma | A2383-5G |
| Lipopolysaccharide from Escherichia coli 0111:B4 (LPS) | Cedarlane | TLRL-3PELPS |
| Human IFN gamma Recombinant Protein | ThermoFisher | PHC4031 |
| Protease inhibitor | Sigma | 11836170001 |
| IL6 ELISA kit | R&D | DY206-05 |
| IL8 ELISA kit | R&D | DY20805 |
| RNeasy MicroKit | Qiagen | 74004 |
| SuperScriptTM VILO™ Mastermix | ThermoFisher | 11755250 |
| PowerUpTM SYBR™ Green Master Mix | ThermoFisher | A25741 |

**Table S2. List of antibodies**

| **Antibody** | **Company** | **Species** | **Dilution** | **Reference number** |
|---|---|---|---|---|
| Cy3-conjugated AffiniPure Donkey Anti-Rabbit IgG | Cedarlane | Donkey | 1: 200 | 711-165-152 |
| Cy5-conjugated AffiniPure Donkey Anti-Mouse IgG | Cedarlane | Donkey | 1: 200 | 715-175-150 |
| Cy5-conjugated AffiniPure Donkey Anti-Rabbit IgG | Cedarlane | Donkey | 1: 200 | 711,175,152 |
| γH2AX | Milipore | Mouse | 1 : 500 | 05-636 |
| H3K9me3 | Abcam | Rabbit | 1: 2000 | AB-176916 |
| p16 | Thermo | Mouse | 1 : 200 | MA517054 |
| p21 | Abcam | Rabbit | 1 : 500 | AB-51243 |
| pHH3 | Cell signaling | Mouse | 1 : 400 | 9706S |
| pRPS6 | Cell signaling | Rabbit | 1: 800 | 4858S |
| HMGB1 | Abcam | Mouse | 1 : 1000 | AB-190377 |
| Lamin B1 | Abcam | Rabbit | 1 : 500 | AB-16048 |

**Table S3. List of primers**

| Gene | Primer | Sequence |
|---|---|---|
| ACTIN | Forward | CCTTGCACATGCCGGAG |
| | Reverse | GCACAGAGCCTCGCCTT |
| GAPDH | Forward | TTGAGGTCAATGAAGGGGTC |
| | Reverse | GAAGGTGAAGGTCGGAGTCA |
| IL-1β | Forward | GAAGCTGATGGCCCTAAACA |
| | Reverse | AAGCCCTTGCTGTAGTGGTG |
| IL-6 | Forward | AGACAGCCACTCACCTCTTCAG |
| | Reverse | TTCTGCCAGTGCCTCTTTGCTG |
| IL8 | Forward | GAGAGTGATTGAGAGTGGACCAC |
| | Reverse | CACAACCCTCTGCACCCAGTTT |
| TNFα | Forward | CTGCTGCACTTTGGAGTGAT |
| | Reverse | AGATGATCTGACTGCCTGGG |

**Table S4. Top 50 enriched GO terms for upregulated genes in SLO vs. VEH experiments.**

| Rank | GO term | Term ID | Adjusted P-val. | Intersection size |
|---|---|---|---|---|
| 1 | regulation of cell migration | GO:0030334 | 1.72E-23 | 133 |
| 2 | regulation of cell motility | GO:2000145 | 2.67E-23 | 138 |
| 3 | cell-cell adhesion | GO:0098609 | 1.76E-18 | 124 |
| 4 | external encapsulating structure organization | GO:0045229 | 4.51E-17 | 64 |
| 5 | cellular response to cytokine stimulus | GO:0071345 | 4.90E-17 | 112 |
| 6 | response to cytokine | GO:0034097 | 1.11E-16 | 119 |
| 7 | extracellular matrix organization | GO:0030198 | 1.51E-16 | 63 |
| 8 | extracellular structure organization | GO:0043062 | 1.79E-16 | 63 |
| 9 | blood vessel development | GO:0001568 | 4.19E-16 | 100 |
| 10 | vasculature development | GO:0001944 | 8.98E-16 | 102 |
| 11 | positive regulation of locomotion | GO:0040017 | 3.50E-15 | 86 |
| 12 | negative regulation of cell population proliferation | GO:0008285 | 5.59E-15 | 96 |
| 13 | positive regulation of cell motility | GO:2000147 | 7.65E-15 | 84 |
| 14 | positive regulation of cell migration | GO:0030335 | 1.34E-14 | 81 |
| 15 | tube morphogenesis | GO:0035239 | 2.31E-14 | 109 |
| 16 | regulation of cell adhesion | GO:0030155 | 8.74E-14 | 100 |
| 17 | negative regulation of developmental process | GO:0051093 | 1.07E-13 | 112 |
| 18 | blood vessel morphogenesis | GO:0048514 | 2.26E-11 | 82 |
| 19 | response to lipid | GO:0033993 | 4.80E-11 | 105 |
| 20 | regulation of cell development | GO:0060284 | 7.42E-11 | 99 |
| 21 | muscle system process | GO:0003012 | 1.99E-10 | 63 |
| 22 | circulatory system process | GO:0003013 | 2.44E-10 | 77 |
| 23 | cytokine-mediated signalling pathway | GO:0019221 | 2.56E-10 | 69 |
| 24 | muscle structure development | GO:0061061 | 2.74E-10 | 84 |
| 25 | negative regulation of locomotion | GO:0040013 | 3.09E-10 | 54 |
| 26 | response to bacterium | GO:0009617 | 4.06E-10 | 87 |
| 27 | negative regulation of cell differentiation | GO:0045596 | 9.48E-10 | 83 |
| 28 | positive regulation of cell differentiation | GO:0045597 | 1.38E-09 | 97 |
| 29 | negative regulation of immune system process | GO:0002683 | 1.63E-09 | 67 |
| 30 | regulation of nervous system development | GO:0051960 | 2.09E-09 | 64 |
| 31 | multicellular organismal-level homeostasis | GO:0048871 | 2.21E-09 | 92 |
| 32 | negative regulation of cell motility | GO:2000146 | 2.62E-09 | 49 |
| 33 | MAPK cascade | GO:0000165 | 2.75E-09 | 87 |
| 34 | blood circulation | GO:0008015 | 4.98E-09 | 67 |
| 35 | cytokine production | GO:0001816 | 5.07E-09 | 89 |
| 36 | regulation of MAPK cascade | GO:0043408 | 6.37E-09 | 78 |
| 37 | cellular anatomical entity morphogenesis | GO:0032989 | 7.75E-09 | 89 |
| 38 | muscle contraction | GO:0006936 | 8.00E-09 | 52 |
| 39 | sensory organ development | GO:0007423 | 8.82E-09 | 75 |
| 40 | tissue morphogenesis | GO:0048729 | 9.56E-09 | 75 |
| 41 | positive regulation of MAPK cascade | GO:0043410 | 1.59E-08 | 62 |
| 42 | response to wounding | GO:0009611 | 1.72E-08 | 70 |
| 43 | regulation of cytokine production | GO:0001817 | 2.05E-08 | 87 |
| 44 | cell morphogenesis | GO:0000902 | 2.12E-08 | 103 |

| | | | | |
|---|---|---|---|---|
| 45 | regulation of cell-cell adhesion | GO:0022407 | 2.37E-08 | 63 |
| 46 | supramolecular fibre organization | GO:0097435 | 2.60E-08 | 92 |
| 47 | angiogenesis | GO:0001525 | 2.67E-08 | 68 |
| 48 | epithelial cell differentiation | GO:0030855 | 3.07E-08 | 84 |
| 49 | ossification | GO:0001503 | 3.51E-08 | 58 |
| 50 | trans-synaptic signalling | GO:0099537 | 3.80E-08 | 85 |

**Table S5. Top 50 enriched GO terms for downregulated genes in SLO vs. VEH experiments.**

| Rank | GO term | Term ID | Adjusted P-val. | Intersection size |
|---|---|---|---|---|
| 1 | translation | GO:0006412 | 1.38E-85 | 363 |
| 2 | peptide biosynthetic process | GO:0043043 | 4.69E-80 | 364 |
| 3 | amide biosynthetic process | GO:0043604 | 1.93E-74 | 396 |
| 4 | ribosome biogenesis | GO:0042254 | 5.17E-68 | 202 |
| 5 | peptide metabolic process | GO:0006518 | 6.63E-67 | 390 |
| 6 | ncRNA metabolic process | GO:0034660 | 2.65E-64 | 305 |
| 7 | chromosome organization | GO:0051276 | 2.43E-58 | 294 |
| 8 | mitotic cell cycle | GO:0000278 | 6.94E-58 | 369 |
| 9 | mitotic cell cycle process | GO:1903047 | 1.02E-56 | 324 |
| 10 | DNA damage response | GO:0006974 | 1.12E-56 | 367 |
| 11 | protein-DNA complex organization | GO:0071824 | 2.11E-51 | 362 |
| 12 | DNA repair | GO:0006281 | 2.81E-51 | 273 |
| 13 | ncRNA processing | GO:0034470 | 4.88E-51 | 221 |
| 14 | chromatin organization | GO:0006325 | 5.60E-45 | 322 |
| 15 | chromosome segregation | GO:0007059 | 1.83E-44 | 207 |
| 16 | regulation of DNA metabolic process | GO:0051052 | 4.32E-44 | 236 |
| 17 | regulation of cell cycle process | GO:0010564 | 3.54E-43 | 293 |
| 18 | cell division | GO:0051301 | 1.37E-39 | 267 |
| 19 | chromatin remodelling | GO:0006338 | 6.02E-38 | 266 |
| 20 | cell cycle phase transition | GO:0044770 | 3.04E-35 | 225 |
| 21 | nuclear chromosome segregation | GO:0098813 | 2.30E-33 | 156 |
| 22 | regulation of translation | GO:0006417 | 2.55E-32 | 173 |
| 23 | mitotic cell cycle phase transition | GO:0044772 | 3.18E-32 | 189 |
| 24 | regulation of cell cycle phase transition | GO:1901987 | 1.69E-31 | 188 |
| 25 | double-strand break repair | GO:0006302 | 1.10E-30 | 149 |
| 26 | positive regulation of DNA metabolic process | GO:0051054 | 6.06E-30 | 145 |
| 27 | regulation of mitotic cell cycle | GO:0007346 | 7.91E-30 | 203 |
| 28 | regulation of mitotic cell cycle phase transition | GO:1901990 | 3.88E-28 | 152 |
| 29 | regulation of mRNA metabolic process | GO:1903311 | 4.76E-27 | 141 |
| 30 | regulation of amide metabolic process | GO:0034248 | 8.33E-27 | 181 |
| 31 | non-membrane-bounded organelle assembly | GO:0140694 | 1.39E-26 | 172 |
| 32 | RNA catabolic process | GO:0006401 | 1.58E-24 | 138 |
| 33 | regulation of cellular response to stress | GO:0080135 | 8.88E-24 | 195 |
| 34 | nuclear transport | GO:0051169 | 3.25E-23 | 144 |
| 35 | nucleocytoplasmic transport | GO:0006913 | 3.25E-23 | 144 |
| 36 | nuclear division | GO:0000280 | 3.50E-23 | 176 |
| 37 | organelle fission | GO:0048285 | 3.95E-23 | 189 |
| 38 | modification-dependent macromolecule catabolic process | GO:0043632 | 3.18E-20 | 215 |
| 39 | nucleobase-containing compound catabolic process | GO:0034655 | 4.41E-19 | 188 |
| 40 | negative regulation of cell cycle process | GO:0010948 | 4.59E-19 | 127 |
| 41 | proteolysis involved in protein catabolic process | GO:0051603 | 6.97E-19 | 239 |

| | | | | |
|---|---|---|---|---|
| 42 | protein modification by small protein conjugation or removal | GO:0070647 | 2.63E-18 | 295 |
| 43 | protein catabolic process | GO:0030163 | 3.86E-18 | 294 |
| 44 | modification-dependent protein catabolic process | GO:0019941 | 8.29E-18 | 206 |
| 45 | intracellular protein transport | GO:0006886 | 9.58E-18 | 225 |
| 46 | DNA recombination | GO:0006310 | 1.66E-17 | 133 |
| 47 | negative regulation of transcription by RNA polymerase II | GO:0000122 | 1.90E-17 | 296 |
| 48 | ubiquitin-dependent protein catabolic process | GO:0006511 | 3.04E-17 | 202 |
| 49 | proteasomal protein catabolic process | GO:0010498 | 6.25E-17 | 175 |
| 50 | heterocycle catabolic process | GO:0046700 | 2.09E-16 | 192 |

**Table S6. Top 50 enriched GO terms for upregulated genes in SLO ½ vs. VEH experiments.**

| Rank | GO term | Term ID | Adjusted P-val. | Intersection size |
|---|---|---|---|---|
| 1 | response to cytokine | GO:0034097 | 2.36E-20 | 92 |
| 2 | cellular response to cytokine stimulus | GO:0071345 | 1.03E-18 | 84 |
| 3 | regulation of cell migration | GO:0030334 | 5.76E-18 | 88 |
| 4 | innate immune response | GO:0045087 | 1.39E-17 | 89 |
| 5 | regulation of cell motility | GO:2000145 | 3.22E-17 | 90 |
| 6 | cytokine-mediated signalling pathway | GO:0019221 | 4.19E-15 | 58 |
| 7 | cell-cell adhesion | GO:0098609 | 7.21E-15 | 83 |
| 8 | response to virus | GO:0009615 | 1.99E-14 | 51 |
| 9 | negative regulation of cell population proliferation | GO:0008285 | 2.43E-13 | 67 |
| 10 | defence response to virus | GO:0051607 | 2.68E-13 | 43 |
| 11 | vasculature development | GO:0001944 | 8.02E-12 | 67 |
| 12 | blood vessel development | GO:0001568 | 1.22E-11 | 65 |
| 13 | external encapsulating structure organization | GO:0045229 | 3.43E-11 | 41 |
| 14 | tube morphogenesis | GO:0035239 | 3.47E-11 | 72 |
| 15 | response to bacterium | GO:0009617 | 1.30E-10 | 63 |
| 16 | positive regulation of response to external stimulus | GO:0032103 | 1.32E-10 | 57 |
| 17 | extracellular matrix organization | GO:0030198 | 1.35E-10 | 40 |
| 18 | regulation of cell adhesion | GO:0030155 | 1.35E-10 | 66 |
| 19 | extracellular structure organization | GO:0043062 | 1.49E-10 | 40 |
| 20 | blood vessel morphogenesis | GO:0048514 | 3.95E-10 | 57 |
| 21 | positive regulation of locomotion | GO:0040017 | 9.56E-10 | 54 |
| 22 | positive regulation of cell motility | GO:2000147 | 1.19E-09 | 53 |
| 23 | positive regulation of cell migration | GO:0030335 | 2.11E-09 | 51 |
| 24 | regulation of cell development | GO:0060284 | 2.51E-09 | 67 |
| 25 | cytokine production | GO:0001816 | 4.33E-09 | 63 |
| 26 | regulation of defence response | GO:0031347 | 7.08E-09 | 62 |
| 27 | regulation of response to biotic stimulus | GO:0002831 | 2.31E-08 | 47 |
| 28 | regulation of cytokine production | GO:0001817 | 2.78E-08 | 61 |
| 29 | regulation of immune response | GO:0050776 | 3.76E-08 | 66 |
| 30 | viral life cycle | GO:0019058 | 3.85E-08 | 36 |
| 31 | regulation of MAPK cascade | GO:0043408 | 4.37E-08 | 54 |
| 32 | angiogenesis | GO:0001525 | 7.56E-08 | 48 |
| 33 | positive regulation of cell population proliferation | GO:0008284 | 8.84E-08 | 69 |
| 34 | regulation of innate immune response | GO:0045088 | 9.55E-08 | 41 |
| 35 | multicellular organismal-level homeostasis | GO:0048871 | 1.54E-07 | 61 |
| 36 | MAPK cascade | GO:0000165 | 1.56E-07 | 58 |
| 37 | negative regulation of immune system process | GO:0002683 | 1.72E-07 | 45 |
| 38 | response to wounding | GO:0009611 | 2.51E-07 | 48 |
| 39 | viral process | GO:0016032 | 2.67E-07 | 41 |
| 40 | muscle structure development | GO:0061061 | 3.18E-07 | 54 |
| 41 | inflammatory response | GO:0006954 | 4.76E-07 | 61 |
| 42 | positive regulation of MAPK cascade | GO:0043410 | 6.99E-07 | 42 |
| 43 | leukocyte activation | GO:0045321 | 7.77E-07 | 66 |
| 44 | regulation of nervous system development | GO:0051960 | 7.99E-07 | 42 |

| | | | | |
|---|---|---|---|---|
| 45 | positive regulation of immune response | GO:0050778 | 9.05E-07 | 55 |
| 46 | positive regulation of transport | GO:0051050 | 9.67E-07 | 61 |
| 47 | ERK1 and ERK2 cascade | GO:0070371 | 1.75E-06 | 33 |
| 48 | negative regulation of response to external stimulus | GO:0032102 | 2.18E-06 | 38 |
| 49 | negative regulation of locomotion | GO:0040013 | 2.41E-06 | 34 |
| 50 | wound healing | GO:0042060 | 2.48E-06 | 39 |

**Table S7. Top 50 enriched GO terms for downregulated genes in SLO ½ vs. VEH experiments.**

| Rank | GO term | Term ID | Adjusted P-val. | Intersection size |
|---|---|---|---|---|
| 1 | translation | GO:0006412 | 1.93E-60 | 289 |
| 2 | protein-DNA complex organization | GO:0071824 | 1.10E-59 | 335 |
| 3 | mitotic cell cycle process | GO:1903047 | 7.01E-59 | 292 |
| 4 | mitotic cell cycle | GO:0000278 | 1.89E-58 | 328 |
| 5 | peptide biosynthetic process | GO:0043043 | 7.87E-58 | 292 |
| 6 | amide biosynthetic process | GO:0043604 | 3.52E-56 | 322 |
| 7 | DNA damage response | GO:0006974 | 4.21E-56 | 324 |
| 8 | chromatin organization | GO:0006325 | 1.17E-55 | 304 |
| 9 | DNA repair | GO:0006281 | 2.76E-53 | 247 |
| 10 | chromosome organization | GO:0051276 | 4.49E-52 | 253 |
| 11 | peptide metabolic process | GO:0006518 | 2.23E-47 | 311 |
| 12 | regulation of cell cycle process | GO:0010564 | 5.21E-43 | 259 |
| 13 | chromatin remodelling | GO:0006338 | 4.10E-42 | 243 |
| 14 | chromosome segregation | GO:0007059 | 1.65E-40 | 180 |
| 15 | regulation of DNA metabolic process | GO:0051052 | 4.75E-37 | 199 |
| 16 | cell division | GO:0051301 | 1.12E-35 | 229 |
| 17 | cell cycle phase transition | GO:0044770 | 3.48E-33 | 196 |
| 18 | regulation of translation | GO:0006417 | 2.87E-31 | 153 |
| 19 | double-strand break repair | GO:0006302 | 1.68E-29 | 132 |
| 20 | mitotic cell cycle phase transition | GO:0044772 | 1.45E-28 | 162 |
| 21 | nuclear chromosome segregation | GO:0098813 | 5.72E-28 | 132 |
| 22 | ncRNA metabolic process | GO:0034660 | 1.21E-27 | 209 |
| 23 | regulation of amide metabolic process | GO:0034248 | 4.51E-27 | 161 |
| 24 | positive regulation of DNA metabolic process | GO:0051054 | 6.29E-26 | 124 |
| 25 | regulation of cell cycle phase transition | GO:1901987 | 1.06E-25 | 157 |
| 26 | microtubule-based process | GO:0007017 | 2.04E-25 | 271 |
| 27 | regulation of mitotic cell cycle | GO:0007346 | 3.25E-25 | 171 |
| 28 | non-membrane-bounded organelle assembly | GO:0140694 | 6.77E-22 | 144 |
| 29 | microtubule cytoskeleton organization | GO:0000226 | 8.78E-22 | 200 |
| 30 | organelle fission | GO:0048285 | 5.68E-21 | 162 |
| 31 | regulation of mRNA metabolic process | GO:1903311 | 7.75E-21 | 116 |
| 32 | regulation of mitotic cell cycle phase transition | GO:1901990 | 1.36E-20 | 123 |
| 33 | nuclear division | GO:0000280 | 1.80E-20 | 150 |
| 34 | nuclear transport | GO:0051169 | 2.50E-20 | 123 |
| 35 | nucleocytoplasmic transport | GO:0006913 | 2.50E-20 | 123 |
| 36 | ribosome biogenesis | GO:0042254 | 2.70E-20 | 120 |
| 37 | regulation of cellular response to stress | GO:0080135 | 1.10E-19 | 163 |
| 38 | negative regulation of transcription by RNA polymerase II | GO:0000122 | 4.97E-19 | 260 |
| 39 | DNA recombination | GO:0006310 | 3.32E-18 | 119 |
| 40 | negative regulation of cell cycle process | GO:0010948 | 5.41E-17 | 109 |
| 41 | positive regulation of cell cycle | GO:0045787 | 6.49E-17 | 120 |
| 42 | protein modification by small protein conjugation or removal | GO:0070647 | 3.99E-16 | 248 |

| | | | | |
|---|---|---|---|---|
| 43 | modification-dependent macromolecule catabolic process | GO:0043632 | 7.05E-16 | 177 |
| 44 | ncRNA processing | GO:0034470 | 7.05E-16 | 138 |
| 45 | RNA catabolic process | GO:0006401 | 1.82E-15 | 107 |
| 46 | positive regulation of organelle organization | GO:0010638 | 8.45E-15 | 150 |
| 47 | intracellular protein transport | GO:0006886 | 3.19E-14 | 186 |
| 48 | negative regulation of cell cycle | GO:0045786 | 3.73E-14 | 123 |
| 49 | ubiquitin-dependent protein catabolic process | GO:0006511 | 9.05E-14 | 167 |
| 50 | modification-dependent protein catabolic process | GO:0019941 | 9.46E-14 | 169 |

**Supplementary video legends**

**Supplementary Video S1. Control VEH condition calcium imaging timelapse.** Representative confocal time lapse images of $Ca^{2+}$ levels in HMC3 cells loaded with 1µM $Ca^{2+}$ indicator Fluo-4 AM. Cells were recorded at 1Hz, over the course of 360 sec. The initial 120 frames were used to define baseline activity, calcium dynamics elicited by 100nM ATP administration were then recorded throughout the following 240 frames. Scale bar = 100µm.

**Supplementary Video S2. SLO treatment condition calcium imaging timelapse.** Representative confocal time lapse images of $Ca^{2+}$ levels in HMC3 cells loaded with 1µM $Ca^{2+}$ indicator Fluo-4 AM. Cells were recorded at 1Hz, over the course of 360 sec. The initial 120 frames were used to define baseline activity, calcium dynamics elicited by 100nM ATP administration were then recorded throughout the following 240 frames. Scale bar = 100µm.
