## Supplemental file 1 for "Chemically induced senescence prompts functional changes in human microglia-like cells"

A modular tool to aggregate results from bioinformatics analyses across many samples into a single report.

#### General Statistics

Copy table | Configure columns | Scatter plot | Violin plot | Showing 18/18 rows and 6/9 columns. | Export as CSV

| Sample Name | % Aligned | M Aligned | % Dropped | % Dups | % GC | M Seqs |
|---|---|---|---|---|---|---|
| SLO_1-2_1 | 91.1 % | 19.8 M | | | | |
| SLO_1-2_1_S35_R1_001 | | | 0.0 % | 52.1 % | 46 % | 21.8 M |
| SLO_1-2_2 | 91.1 % | 19.8 M | | | | |
| SLO_1-2_2_S36_R1_001 | | | 0.0 % | 51.7 % | 46 % | 21.8 M |
| SLO_1-2_3 | 91.2 % | 18.6 M | | | | |
| SLO_1-2_3_S37_R1_001 | | | 0.0 % | 51.9 % | 46 % | 20.4 M |
| SLO_1_1 | 91.7 % | 18.2 M | | | | |
| SLO_1_1_S38_R1_001 | | | 0.0 % | 50.3 % | 46 % | 19.8 M |
| SLO_1_2 | 91.9 % | 18.8 M | | | | |
| SLO_1_2_S39_R1_001 | | | 0.0 % | 49.3 % | 46 % | 20.4 M |
| SLO_1_3 | 91.9 % | 16.7 M | | | | |
| SLO_1_3_S40_R1_001 | | | 0.0 % | 48.9 % | 46 % | 18.2 M |
| VEH_1 | 90.7 % | 18.0 M | | | | |
| VEH_1_S32_R1_001 | | | 0.0 % | 53.0 % | 46 % | 19.8 M |
| VEH_2 | 90.7 % | 17.7 M | | | | |
| VEH_2_S33_R1_001 | | | 0.0 % | 51.9 % | 46 % | 19.5 M |
| VEH_3 | 90.5 % | 22.0 M | | | | |
| VEH_3_S34_R1_001 | | | 0.0 % | 55.1 % | 46 % | 24.3 M |

### RSeQC

RSeQC package provides a number of useful modules that can comprehensively evaluate high throughput RNA-seq data. *DOI: 10.1093/bioinformatics/bts356.*

#### Read Distribution

Read Distribution calculates how mapped reads are distributed over genome features.

Percentages | Export Plot

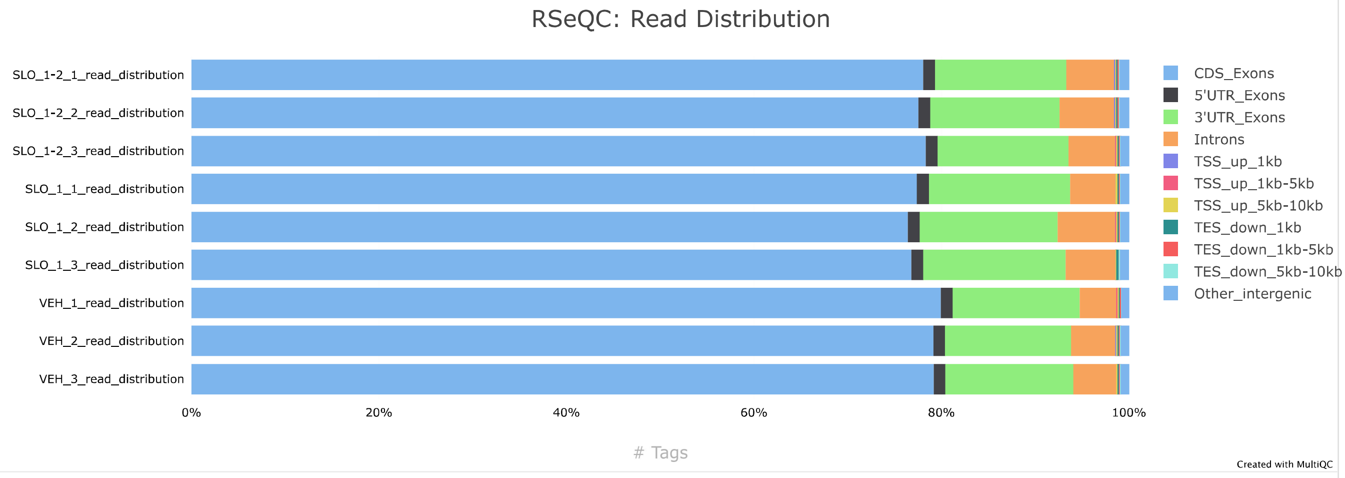

### STAR

STAR is an ultrafast universal RNA-seq aligner. *DOI: 10.1093/bioinformatics/bts635.*

#### Alignment Scores

Percentages

Export Plot

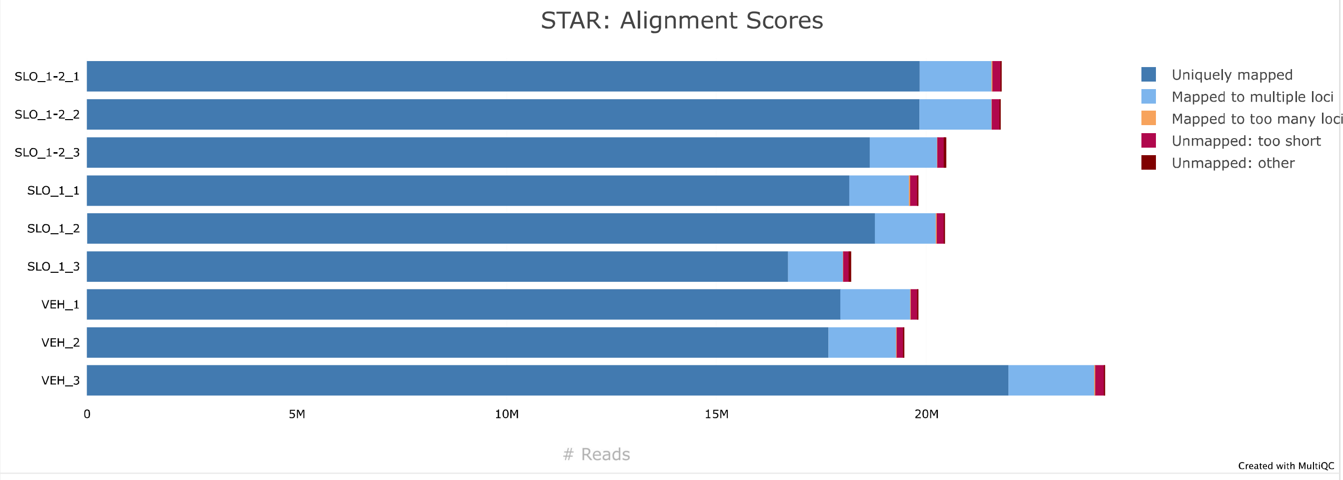


#### Gene Counts

Statistics from results generated using `--quantMode GeneCounts` . The three tabs show counts for unstranded RNA-seq, counts for the 1st read strand aligned with RNA and counts for the 2nd read strand aligned with RNA.

Percentages | Unstranded | Same Stranded | Reverse Stranded

Export Plot

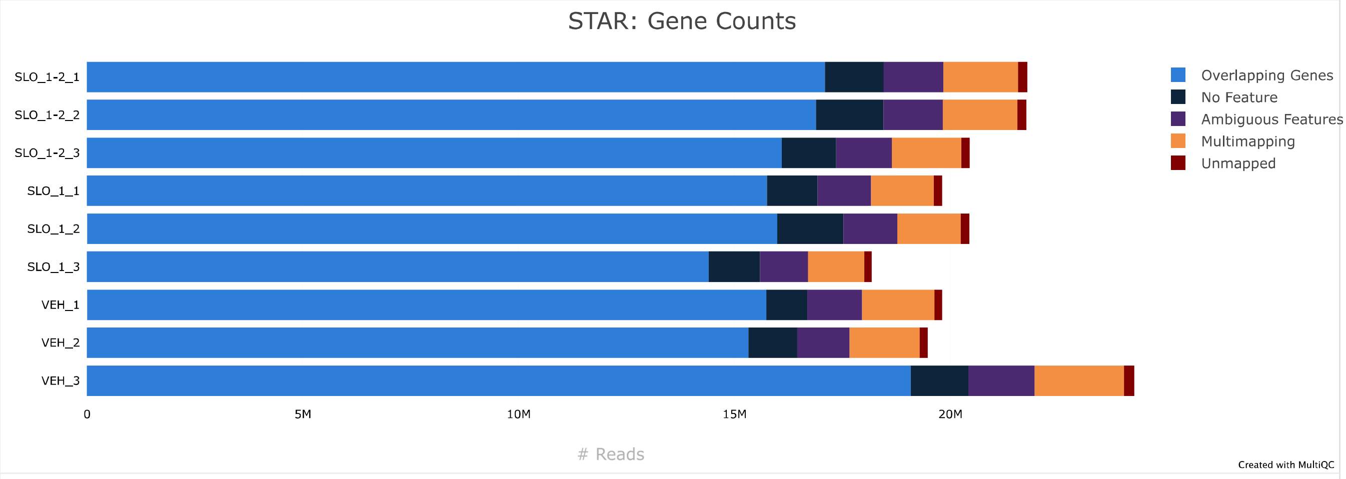


### Trimmomatic

Trimmomatic is a flexible read trimming tool for Illumina NGS data. *DOI: 10.1093/bioinformatics/btu170.*

Percentages

Export Plot

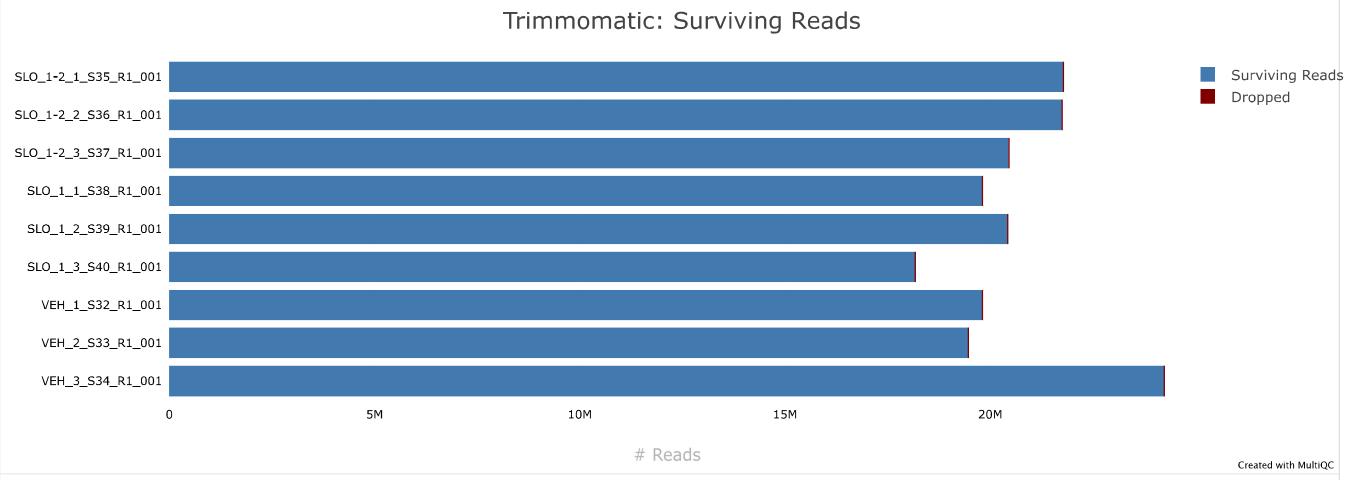

### FastQ Screen *Version:* `0.14.0`

FastQ Screen allows you to screen a library of sequences in FastQ format against a set of sequence databases so you can see if the composition of the library matches with what you expect. *DOI: 10.12688/f1000research.15931.2.*

#### Mapped Reads

Percentages Export Plot

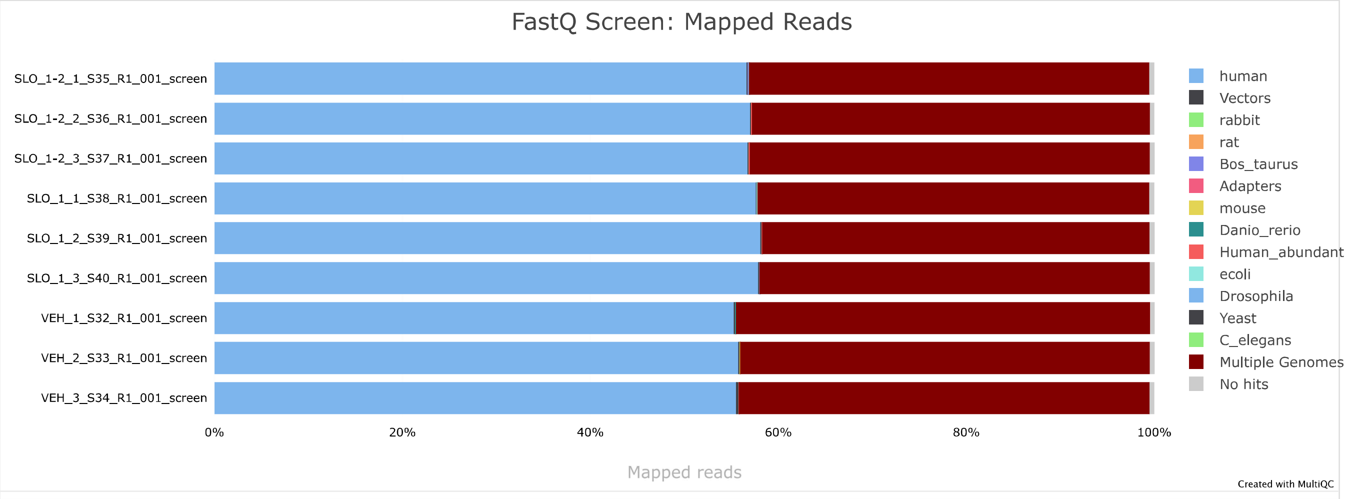


### FastQC *Version:* `0.11.9`

FastQC is a quality control tool for high throughput sequence data, written by Simon Andrews at the Babraham Institute in Cambridge.

#### Sequence Counts

Sequence counts for each sample. Duplicate read counts are an estimate only.

Percentages Export Plot

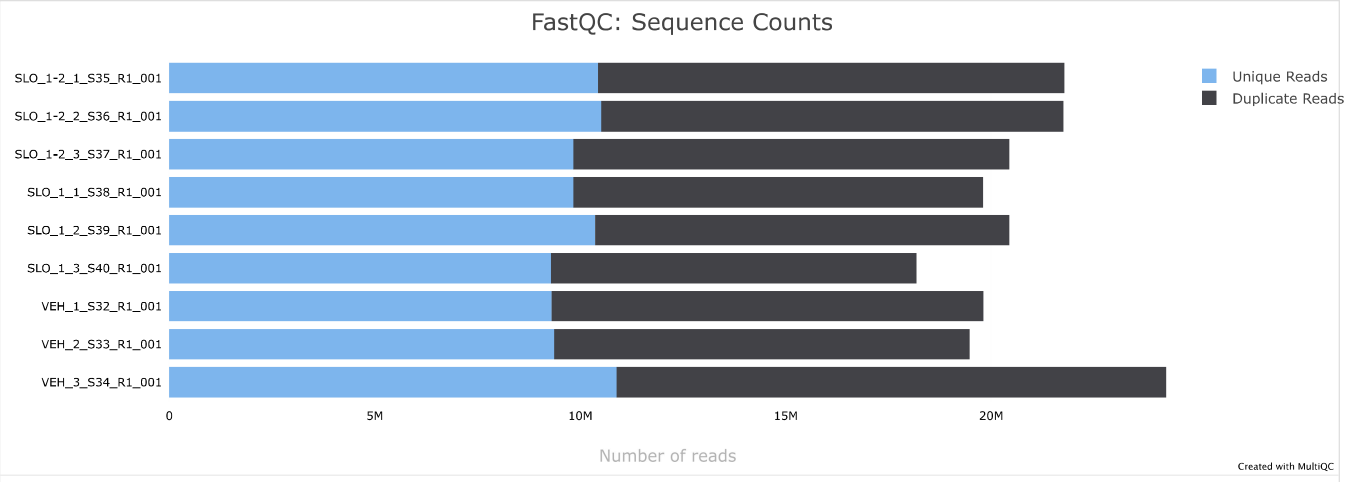

#### Sequence Quality Histograms 9

The mean quality value across each base position in the read.

Export Plot

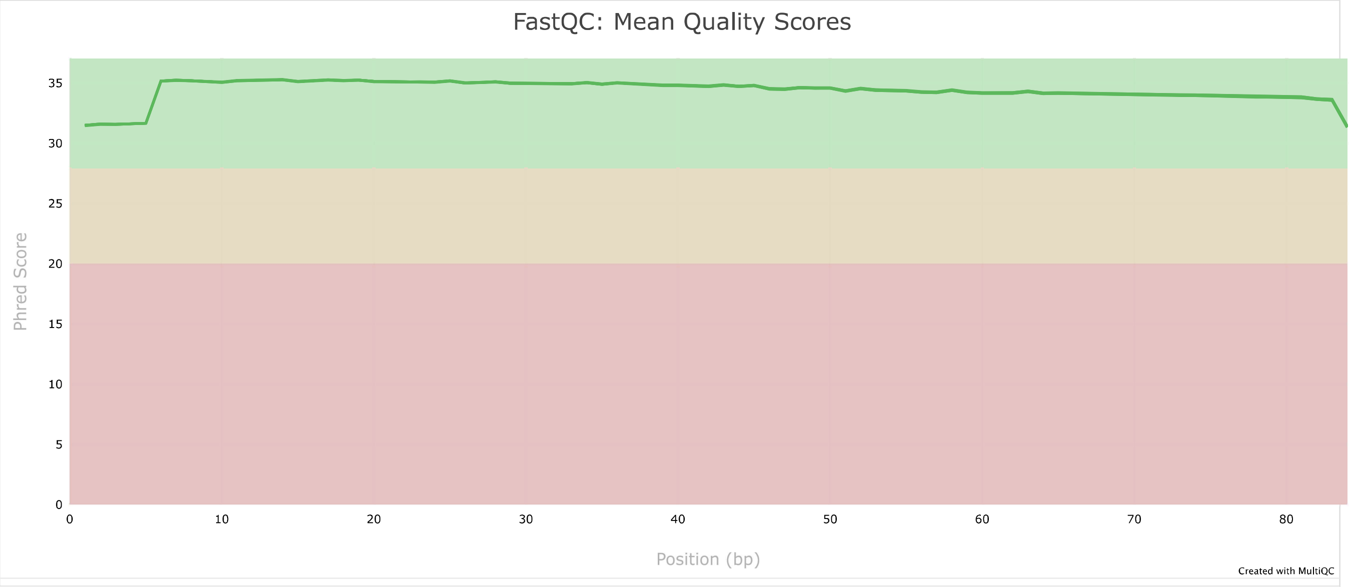


#### Per Sequence Quality Scores 9

The number of reads with average quality scores. Shows if a subset of reads has poor quality.

Export Plot

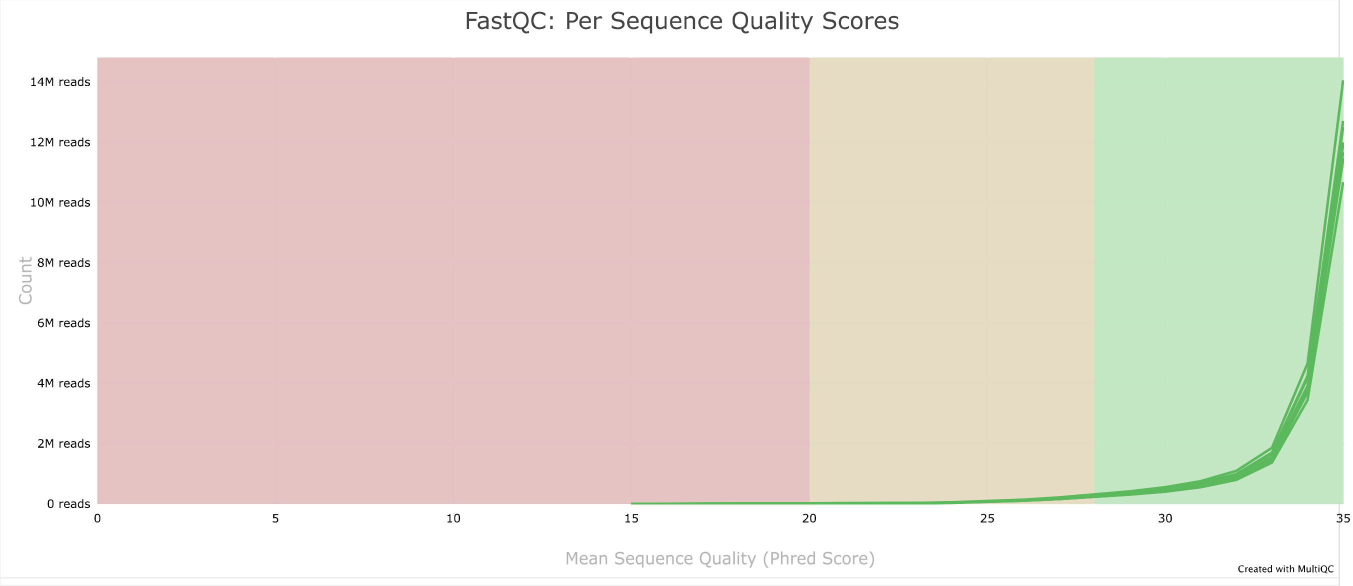

#### Per Base Sequence Content 9

The proportion of each base position for which each of the four normal DNA bases has been called.

Click a sample row to see a line plot for that dataset.

Rollover for sample name

Position: - %T: - %C: - %A: - %G: -

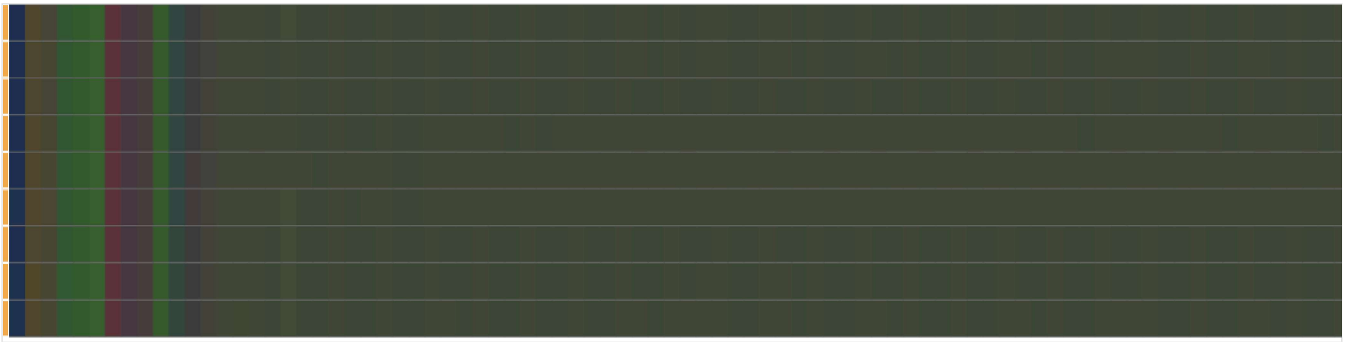

#### Per Sequence GC Content 9

The average GC content of reads. Normal random library typically have a roughly normal distribution of GC content.

Percentages Counts Export Plot

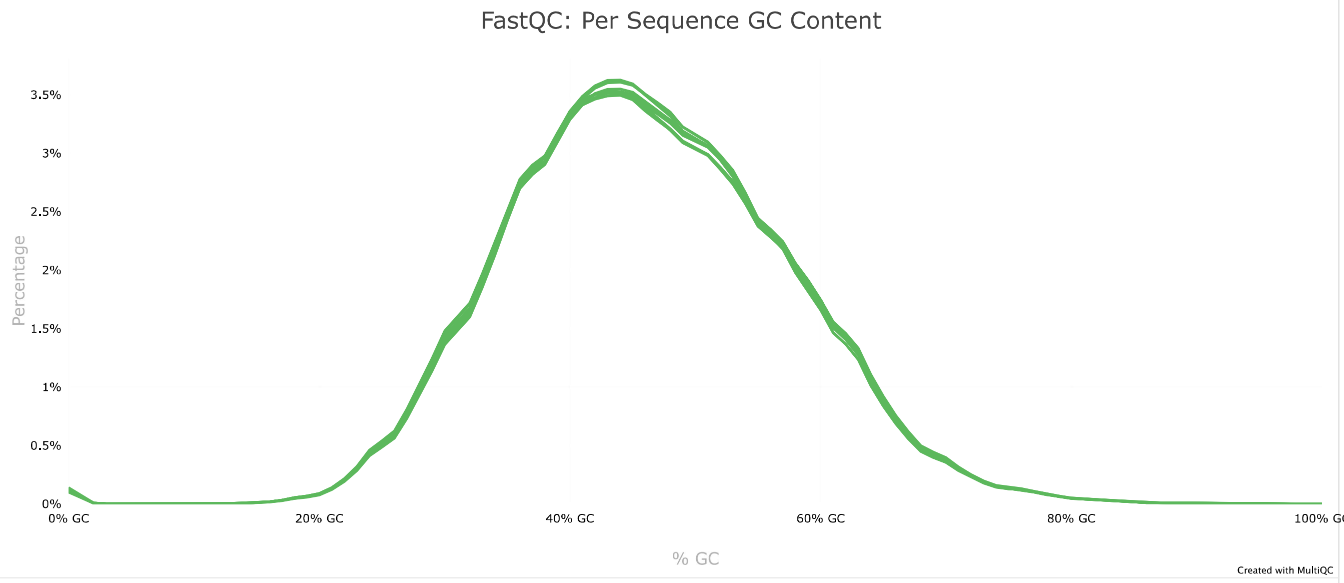

#### Per Base N Content 9

The percentage of base calls at each position for which an N was called.

Export Plot

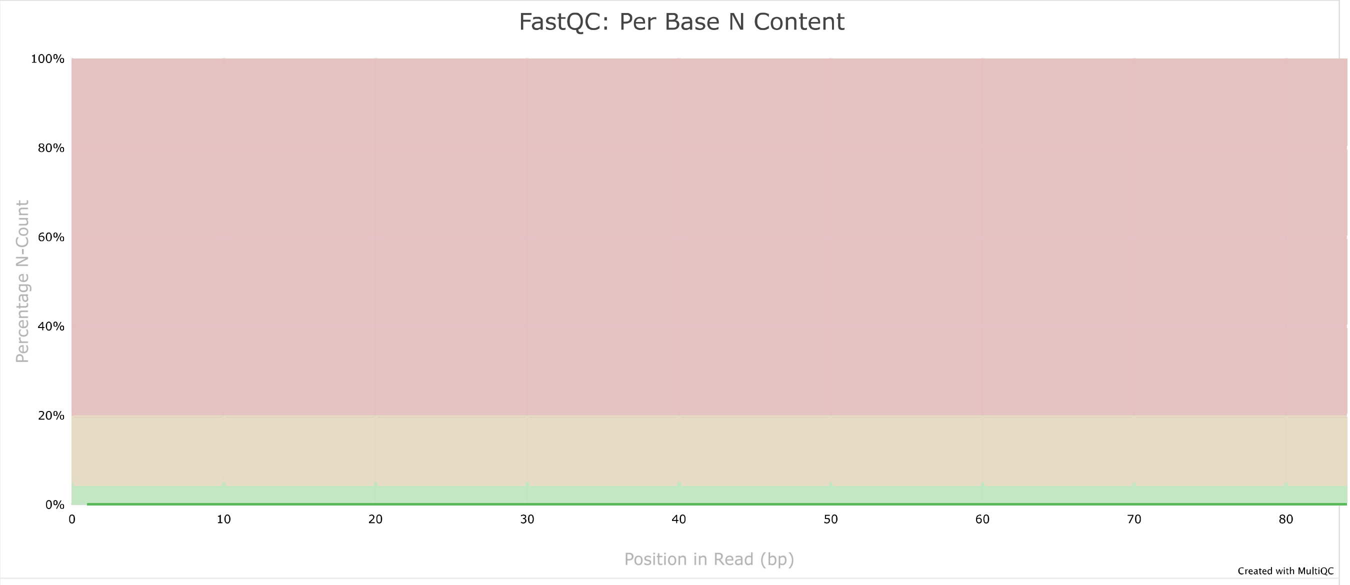


#### Sequence Length Distribution 9

All samples have sequences of a single length (84bp).

#### Sequence Duplication Levels 2

The relative level of duplication found for every sequence.

Export Plot

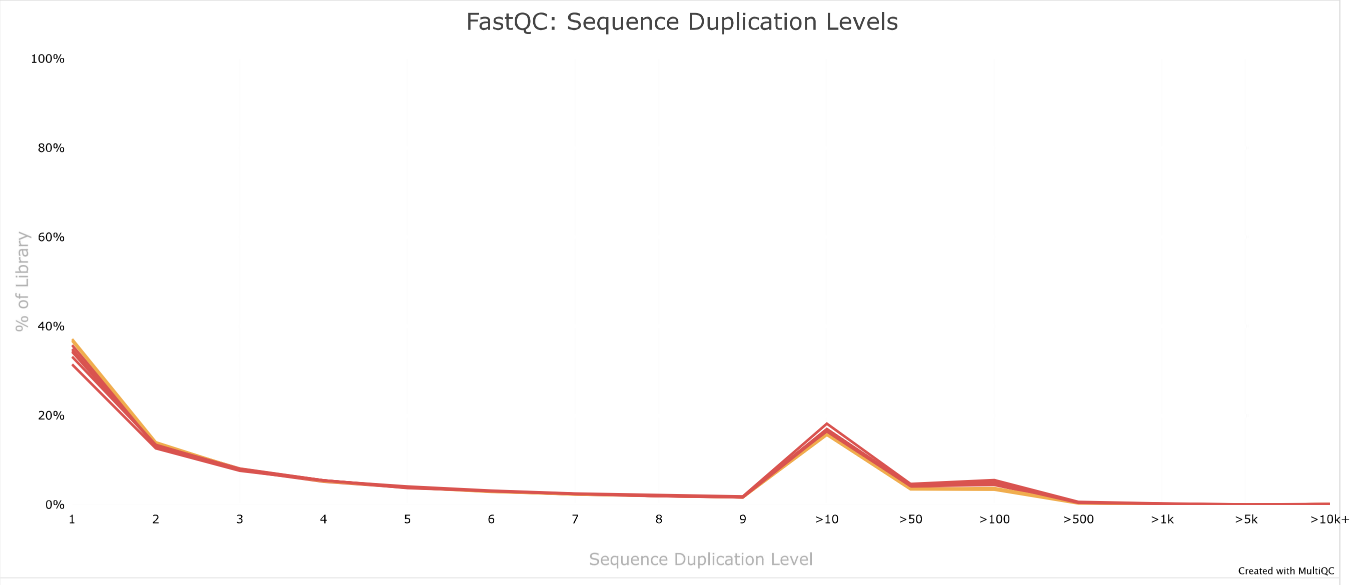


#### Overrepresented sequences by sample 9

The total amount of overrepresented sequences found in each library.

9 samples had less than 1% of reads made up of overrepresented sequences

#### Top overrepresented sequences

Top overrepresented sequences across all samples. The table shows 20 most overrepresented sequences across all samples, ranked by the number of samples they occur in.

Copy table | Configure columns | Scatter plot | Violin plot | Showing 1/1 rows and 3/3 columns. | Export as CSV

| Overrepresented sequence | Samples | Occurrences | % of all reads |
|---|---|---|---|
| TTTTTTTTTTTTTTTTTTTTTTTTTTTT | 9 | 226 467 | 0.1218 % |

#### Adapter Content 9

The cumulative percentage count of the proportion of your library which has seen each of the adapter sequences at each position.

Export Plot

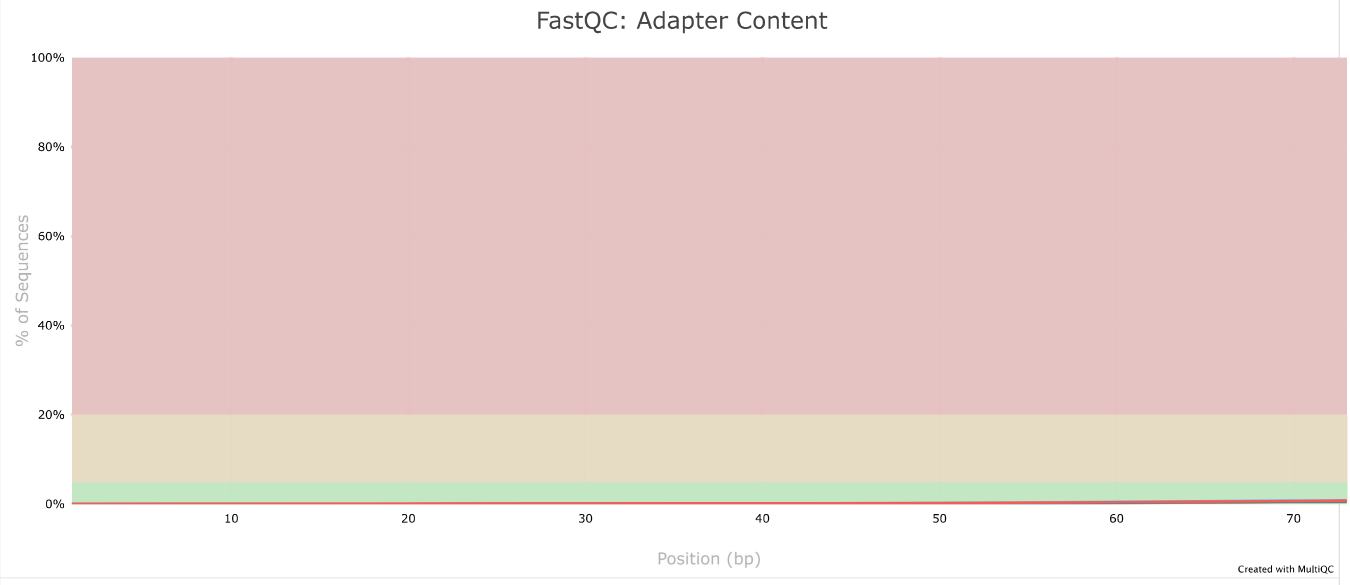


#### Status Checks

Status for each FastQC section showing whether results seem entirely normal (green), slightly abnormal (orange) or very unusual (red).

**Min:** 0 **Max:** 1 Export Plot

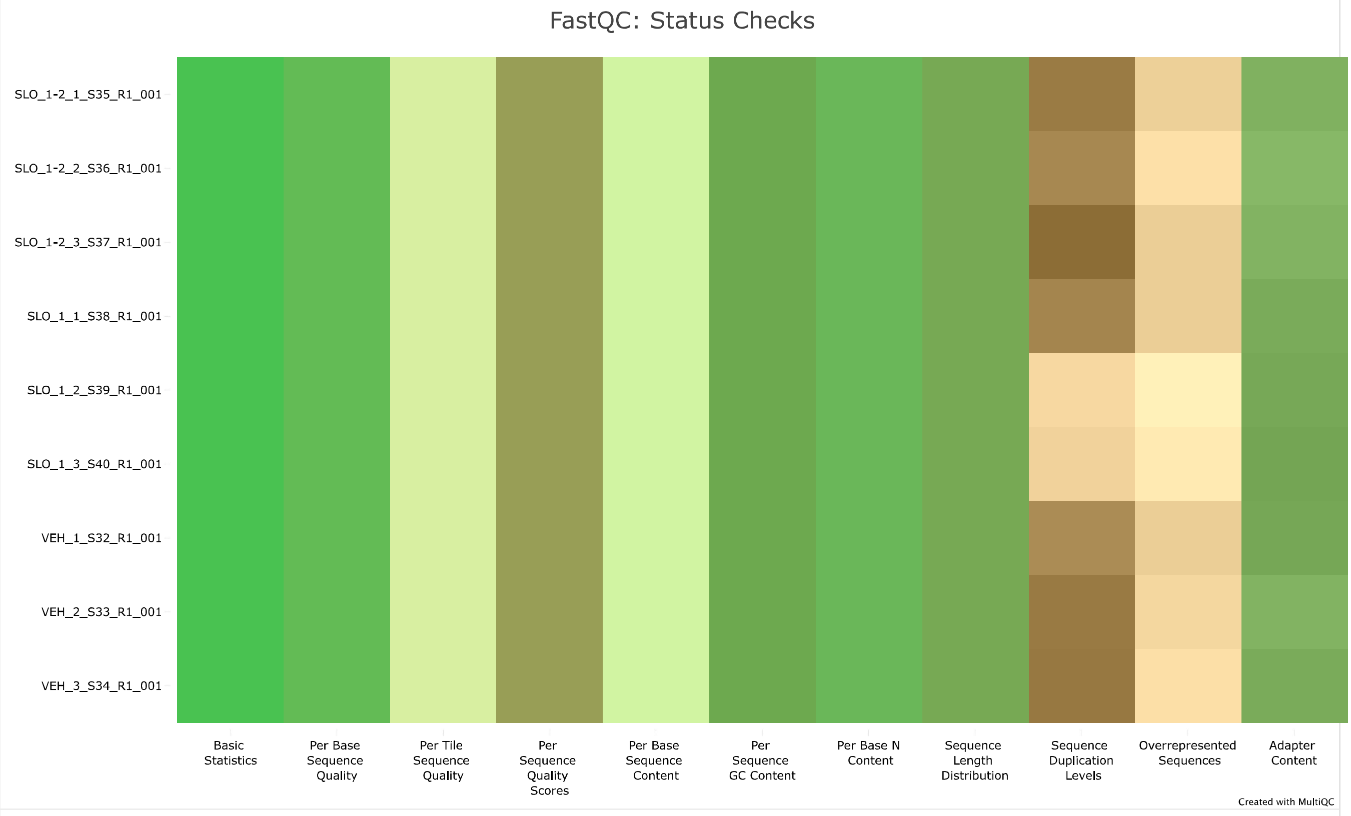

### Software Versions

Software Versions lists versions of software tools extracted from file contents.

Copy table

| Software | Version |
|---|---|
| FastQ Screen | 0.14.0 |
| FastQC | 0.11.9 |

**MultiQC v1.21** - Written by Phil Ewels, available on GitHub.
This report uses HighCharts, jQuery, jQuery UI, Bootstrap, FileSaver.js and clipboard.js.

seqera
